# L-Lactate reprograms tumor-associated macrophages to drive pancreatic cancer progression via BCL3 lactylation

**DOI:** 10.64898/2026.03.06.710237

**Authors:** Pei-Xuan Ji, Jia-Hao Zheng, Xue-Shiyu Ma, Feng Yu, Ya-Wen Lu, Qi Wo, Kan Xu, Min-Wei Yang, Jian-Yu Yang, Wei Liu, Xue-Liang Fu, Yan-Miao Huo, Wen-Yi Gu, Yu-Jun Hao, Shu-Heng Jiang, De-Jun Liu, Yong-Wei Sun

## Abstract

Metabolic reprogramming fuels pancreatic ductal adenocarcinoma (PDAC) malignancy, creating a nutrient-deprived and waste-rich microenvironment. How this extreme metabolic pressure dictates the phenotypic remodeling of infiltrating immune cells remains largely unclear. Metabolomic profiling of PDAC reveals that robust tumor glycolysis proceeds without a commensurate accumulation of extracellular lactate. Integrated single-cell profiling and tissue multiplex immunofluorescence demonstrate that tumor-associated macrophages (TAMs) act as the primary consumers of tumor-secreted lactate. The uptake of L-lactate directs macrophages toward a pro-tumorigenic state through site-specific L-lactylation of the transcriptional co-regulator BCL3 at lysine 21 (K21). Functioning as a molecular switch, K21 lactylation triggers BCL3 nuclear translocation and enhances its interaction with the NF-κB p50 subunit. The ensuing BCL3-p50 complex competitively displaces the pro-inflammatory p65 subunit, rewiring the transcriptional output to suppress inflammation and enforce tumor-supporting networks. *In vivo*, macrophage-specific expression of a lactylation-deficient BCL3 mutant (K21R) abolishes lactate-driven phenotypic shifts and restricts tumor growth. Clinically, a BCL3-lactylated macrophage signature spatially correlates with CD8^+^ T cell exclusion and predicts poor patient survival, providing a strong rationale for targeting the BCL3-lactylation axis to reverse TAM-driven PDAC progression.

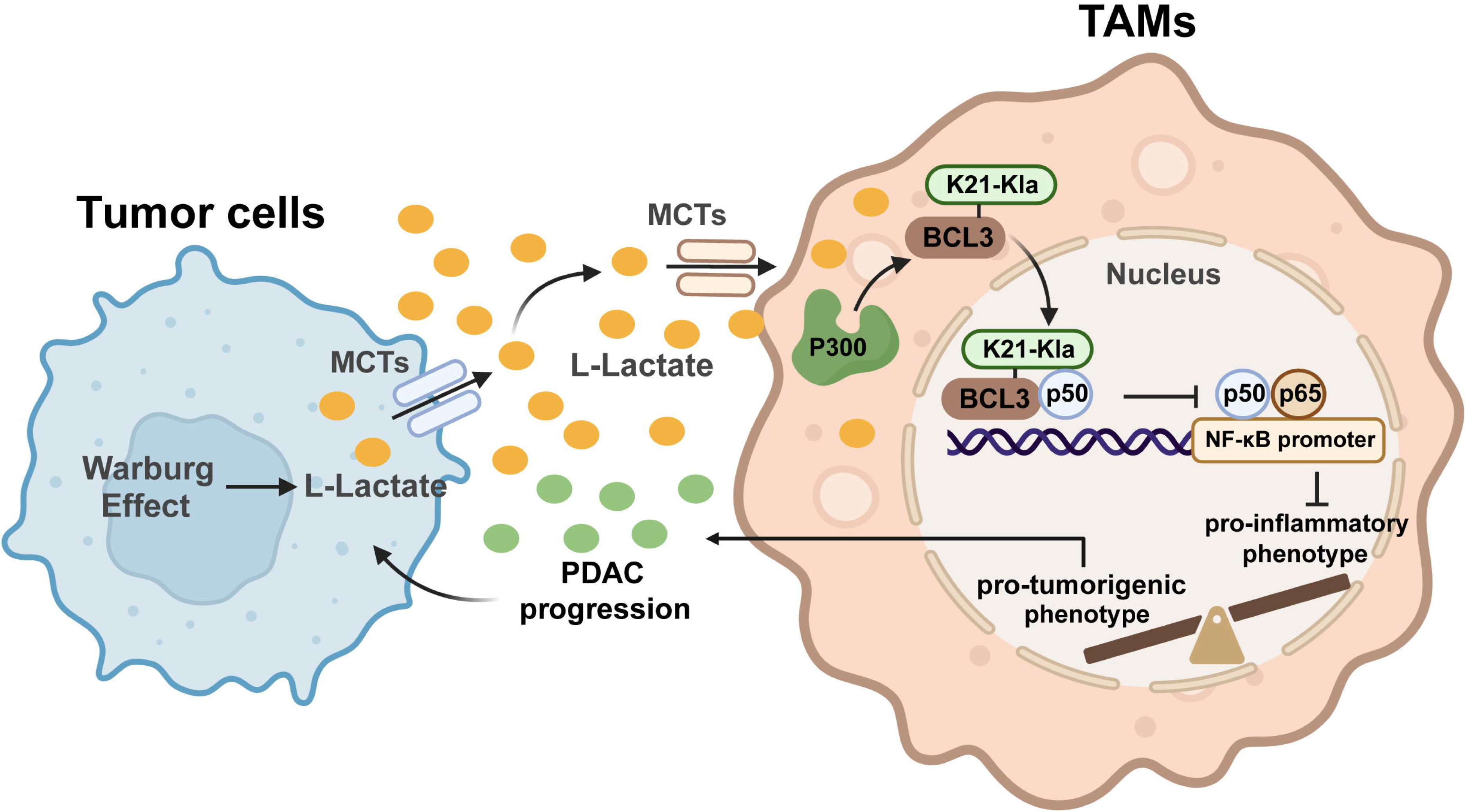

## Introduction

Pancreatic ductal adenocarcinoma (PDAC) features a dense, desmoplastic tumor microenvironment (TME) that functions as a physical and immunosuppressive barrier to therapy.^1-3^ Within this TME, Tumor-associated macrophages (TAMs) constitute the predominant immune population.^4,5^ While extensive evidence shows that PDAC-associated macrophages acquire a pro-tumorigenic M2-like phenotype to promote angiogenesis, fibrosis, and immune evasion, the specific upstream signals driving the functional reprogramming remain unclear.^3,4,6^ Consequently, therapeutic strategies targeting differentiated M2 macrophages have yielded limited success,^7^ highlighting an urgent need to identify the key mechanisms governing macrophage plasticity.

Metabolic reprogramming is a core feature of PDAC, characterized by elevated aerobic glycolysis and significant lactate accumulation.^8,9^ Historically viewed as a metabolic by-product, lactate is now recognized as a pivotal signaling molecule that determines cell fate within the TME.^10,11^ The recent discovery of protein lactylation, a post-translational modification (PTM) derived from L-lactate, provides a direct link between metabolite accumulation and gene regulation.^12,13^ Given the highly glycolytic nature of PDAC, tumor-derived lactate likely functions as an intercellular messenger shaping the local immune profile.^14,15^ However, how specific immune cells exploit extracellular lactate to induce distinct transcriptional programs remains largely unknown.

NF-κB signaling is essential for balancing pro- and anti-inflammatory macrophage responses.^16^ B-cell lymphoma 3 (BCL3), an atypical IκB family member, modulates NF-κB transcriptional activity by directly interacting with its subunits.^17^ While elevated BCL3 expression correlates with poor prognosis in various malignancies^18,19^ and is known to suppress pro-inflammatory cytokines for limiting excessive inflammation in macrophages,^20,21^ the upstream signals regulating its nuclear translocation and interaction dynamics within the TME remain undefined. Specifically, whether BCL3 functions as a “responder” or “mediator” for metabolic alterations, including lactylation, to couple the lactate-rich environment to tumor-supporting gene expression is unexplored.

Here, using integrated metabolomics and single-cell RNA sequencing (scRNA-seq), we identify a metabolic crosstalk in which macrophages preferentially consume PDAC-derived lactate. Our data demonstrate that lactate uptake acts as a direct signaling cue, driving TAMs polarization toward a pro-tumorigenic M2-like phenotype to promote PDAC progression. Mechanistically, lactate induces site-specific L-lactylation of BCL3 at lysine 21 (K21). The resulting modification promotes BCL3 nuclear translocation and stabilizes its interaction with the NF-κB p50 subunit, thereby competitively displacing p65 and repressing downstream pro-inflammatory gene expression. Mapping the metabolic-signaling axis reveals a key mechanism of PDAC progression and identifies BCL3 lactylation as a potential therapeutic target to reverse macrophage-mediated tumor-promoting effects.

## Methods and Materials

### Human Tissue Specimens and Ethical Statement

Human PDAC tissues and paired adjacent non-tumor tissues were prospectively collected from patients undergoing curative surgical resection at the Department of Biliary-Pancreatic Surgery, Ren Ji Hospital. Prior to sample acquisition, written informed consent was obtained from all participants. The study protocol was approved by the Institutional Review Board (IRB) of Ren Ji Hospital (Approval No. RA-2025-251). Fresh surgical specimens were immediately aliquoted: one portion was flash-frozen in liquid nitrogen and stored at -80°C for downstream metabolomic and proteomic analyses, while the corresponding portion was fixed in 4% paraformaldehyde (PFA) and embedded in paraffin for histopathological verification.

### Cell Lines and Cell Culture

The human PDAC cell lines PANC-1 (RRID: CVCL_0480) and PATU8988 (RRID: CVCL_1846), the human monocytic cell line THP-1 (RRID: CVCL_0006), and the murine macrophage cell line RAW 264.7 (RRID: CVCL_0493) were obtained from the National Collection of Authenticated Cell Cultures, Chinese Academy of Sciences (Shanghai, China). The murine PDAC cell lines (KPC1199 and Panc02) and the human patient-derived PDAC cell line (PDC0034) were kindly provided by Professor Jing Xue (Ren Ji Hospital, Shanghai Jiao Tong University). Cells were cultured in DMEM (Procell, Cat# PM150210) or RPMI-1640 (Procell, Cat# PM150110) supplemented with 10% Fetal Bovine Serum (FBS; ExCell Bio, Cat# FSP500) and 1% Penicillin-Streptomycin Solution (Share-bio, Cat# SB-CR011). All cultures were maintained at 37°C in a humidified incubator containing 5% CO_2_. Routine testing was performed to ensure cells were free of mycoplasma contamination.

To generate macrophage-like phenotypes, THP-1 monocytes were differentiated using 100 ng/mL phorbol 12-myristate 13-acetate (PMA; MedChemExpress, Cat# HY-18739) for 24 hours. Primary bone marrow-derived macrophages (BMDMs) were isolated from the femurs and tibias of 6- to 8-week-old C57BL/6J mice and differentiated for 7 days in RPMI-1640 medium containing 20 ng/mL macrophage colony-stimulating factor (M-CSF; MedChemExpress, Cat# HY-P7085).

### Reagents and Antibodies

Chemical reagents, including Sodium L-lactate (L-NaLac, Cat# 71718), Sodium D-lactate (D-NaLac, Cat# 71716), Sodium Acetate (NaAce, Cat# S2889), and Lipopolysaccharides (LPS, Cat# L2880), were purchased from Sigma-Aldrich (St. Louis, MO, USA). Pharmacological inhibitors, including AR-C155858 (MCT1/2 inhibitor, Cat# HY-13248), 7ACC1 (MPC inhibitor, Cat# HY-D0067), C646 (p300 inhibitor, Cat# HY-13823), JS6 (BCL3 inhibitor, Cat# HY-152177), PX-478 (HIF-1α inhibitor, Cat# HY-10231), GSK-2837808A (LDHA inhibitor, Cat# HY-100681), and Lactic Acid (Cat# HY-B2227), were obtained from MedChemExpress (Monmouth Junction, NJ, USA).

For immunoblotting and immunofluorescence assays, high-specificity primary antibodies were utilized. Key antibodies included rabbit monoclonal anti-L-Lactyl Lysine (PTM Biolabs, Cat# PTM-1401RM, RRID: AB_2942013), anti-BCL3 (Zen Bioscience, Cat# 860954), anti-ARG1 (Proteintech Group, Cat# 16001-1-AP, RRID: AB_2289842), anti-CD86 (Cell Signaling Technology, Cat# 91882, RRID: AB_2797422), anti-NF-κB p105/p50 (Genuin Biotech, Cat# 61524), and anti-NF-κB p65 (Proteintech, Cat# 80979-1-RR, RRID: AB_2918923). A complete inventory of antibodies is provided in Supplementary Table S1.

### Lentiviral Transduction

To achieve stable gene knockdown, lentiviral vectors encoding short hairpin RNAs (shRNAs) targeting human *LDHA* and murine *Ldha*, along with a non-targeting scramble control (shNC), were custom-synthesized based on validated target sequences. For murine *Bcl3* knockdown, the OmicsLink™ shRNA Expression Clone (MSH036845-LVRU6GP-a) was obtained from GeneCopoeia, Inc. (Rockville, MD, USA). For functional rescue experiments, wild-type (WT) and lactylation-deficient K21R mutant murine *Bcl3* constructs were purchased as ready-to-use OmicsLink™ Expression Clones (CS-Mm24400-Lv158-01 and CS-Mm24400-Lv158-02, respectively; GeneCopoeia). Lentiviral particles were produced by co-transfecting HEK293T cells with the respective transfer plasmids and packaging vectors (psPAX2 and pMD2.G) using Polybrene (Beyotime, #C0351). Viral supernatants were collected at 48 and 72 hours post-transfection, filtered through a 0.45 μm membrane, and utilized for target cell infection. Stable cell lines were selected using puromycin (2-5 μg/mL) for 7 days. Transduction efficiency was confirmed by RT-qPCR and Western blot analysis. Oligonucleotide sequences and specific clone details are provided in Supplementary Table S2.

### Histopathology and Immunohistochemistry (IHC)

Tissue specimens were fixed in 4% PFA, embedded in paraffin, and sectioned at a thickness of 4 µm. For morphological evaluation, sections underwent standard deparaffinization in xylene and rehydration through a graded ethanol series, followed by hematoxylin and eosin (H&E) staining. For IHC analysis, antigen retrieval was performed by submerging sections in citrate buffer (pH 6.0) and heating via microwave irradiation for 15 minutes, followed by cooling to room temperature. To minimize non-specific background staining, endogenous peroxidase activity was quenched with 0.3% hydrogen peroxide (H_2_O_2_) for 15 minutes, and sections were blocked with 10% Bovine Serum Albumin (BSA; Solarbio, Cat# A8020) for 1 hour at room temperature.

Subsequently, sections were incubated overnight at 4°C with primary antibody against Ki-67 (Proteintech, Cat# 27309-1-AP, RRID: AB_2756525, dilution 1:4000) diluted in blocking buffer. Following three washes with PBS, sections were incubated with HRP-conjugated secondary antibodies (Share-bio, Cat# SB-AB0101/SB-AB0102) for 1 hour at room temperature. Immunoreactivity was visualized using a 3,3’-diaminobenzidine substrate kit (DAB; Proteintech, Cat# PR30018), and nuclei were counterstained with hematoxylin. Images were captured using a standard bright-field microscopy system.

### Multiplex Immunofluorescence (mIF) Staining

Multiplex immunofluorescence was performed on formalin-fixed, paraffin-embedded (FFPE) tissue specimens using the Fluorescein TSA Fluorescence System Kit (Runnerbio, Cat# bry-0034-100). Following deparaffinization in xylene and rehydration through a graded ethanol series, heat-induced epitope retrieval (HIER) was performed in Tris-EDTA buffer (pH 9.0) to expose antigenic sites. Endogenous peroxidase activity was quenched, and non-specific binding sites were blocked with 5% BSA. The staining protocol followed a sequential immunolabeling strategy: each cycle consisted of primary antibody incubation overnight at 4°C, followed by detection with a horseradish peroxidase (HRP)-conjugated secondary antibody and covalent tagging using distinct Tyramide Signal Amplification (TSA) fluorophores. To facilitate multiplexing, the primary antibody-HRP complex was stripped via microwave heating in citrate buffer (pH 6.0) between successive cycles, while the fluorophores remained covalently bound to the antigen. Finally, nuclei were counterstained with DAPI (Beyotime, Cat# C1006), and slides were mounted with an antifade mounting medium.

For quantitative assessment, multispectral images were acquired and processed using the ImageJ/Fiji platform (NIH). Individual channels were spectrally unmixed and thresholded to define single-marker positivity. To ensure data representativeness and mitigate intratumoral heterogeneity, a systematic sampling strategy was employed. For TMAs, the entire viable tissue area of each core was analyzed, excluding necrotic and folded regions. For whole-tissue sections (human and mouse), three to five non-overlapping Regions of Interest (ROIs) were randomly selected within the intratumoral and peritumoral compartments at high magnification (200× or 400×). The ‘Image Calculator’ function utilizing the ‘AND’ boolean operator was applied to isolate pixels exhibiting strict co-localization of CD68, BCL3, and Pan-Kla signals. The density of these triple-positive regions was quantified using the ‘Analyze Particles’ plugin. The final BCL3-Kla Macrophage Score (BKMS) for each specimen was calculated as the mean density of the analyzed ROIs, normalized to the tissue surface area (mm²).

### Co-culture Systems and Conditioned Medium

Non-contact co-cultures were established using Transwell inserts to permit metabolite diffusion while preventing direct cell-cell contact. To evaluate the education of macrophages by tumor cells, PDAC cells (subjected to specific genetic or pharmacological interventions) were seeded in the upper inserts (2 × 10^5^ cells/well), while macrophages were cultured in the lower chambers (4 × 10^5^ cells/well). Conversely, for macrophage migration assays, macrophages were seeded in the upper inserts.

For preparation of conditioned medium (CM), PDAC cells or macrophages were cultured to 70-80% confluence. The cells were then washed with PBS and replenished with fresh medium in the presence or absence of specific stimuli (e.g., lactate, LPS). After 48 hours, supernatants were collected, centrifuged (2000 g, 10 minutes) to eliminate cellular debris, and filtered through a 0.22 μm membrane. The resulting CM was used immediately or stored at -80°C for subsequent treatment of recipient cells to evaluate soluble factor-mediated signaling.

### Transwell Migration Assay

Chemotactic migration was assessed using Transwell chambers with 5.0 μm pore size polycarbonate membranes (Corning, Cat# 3421). Macrophages (1 × 10^5^ cells) suspended in 200 μL serum-free medium were seeded into the upper chamber. The lower chamber was filled with 600 μL of either complete medium or PDAC-CM acting as a chemoattractant. After 24 hours of incubation, non-migrating cells on the upper surface were removed via cotton swabs. Cells that migrated to the lower surface were fixed with 4% PFA (Beyotime, Cat# P0099) for 20 minutes, stained with 0.1% crystal violet for 20 minutes, and quantified by counting five random high-power fields under a light microscope.

### Colony Formation Assay

PDAC cells or macrophages were seeded at a low density (500-1000 cells per well) into 6-well plates. Cells were cultured for 10-14 days, with the medium replenished every 3 days. Upon the formation of visible colonies (> 50 cells), cultures were fixed with 4% PFA for 15 minutes and stained with 0.1% crystal violet. Colonies were imaged and counted using ImageJ software.

### EdU Incorporation Assay

DNA synthesis activity was assessed using the 488 Click-iT 5-ethynyl-2’-deoxyuridine (EdU) Green Cell Proliferation Kit (Share-bio, Cat# SB-C6043). Cells were seeded on sterile glass coverslips or in 24-well plates (1 × 10^5^ cells/well) and subjected to the indicated treatments. Subsequently, cells were incubated with 50 μM EdU medium for 2 hours at 37°C. Following the labeling period, cells were washed with PBS and fixed with 4% paraformaldehyde for 30 minutes at room temperature. To ensure antibody and dye penetration, membranes were permeabilized using 0.5% Triton X-100 in PBS for 20 minutes. The click chemistry reaction was performed by incubating cells with the Apollo fluorescent dye solution for 30 minutes in the dark. Nuclei were counterstained with DAPI (1 μg/mL) for 5 minutes. Images were acquired using a fluorescence microscope. The proliferation rate was quantified by calculating the ratio of EdU-positive nuclei (green fluorescence) to total DAPI-stained nuclei (blue fluorescence) in at least five randomly selected fields per sample using ImageJ software.

### Lactate Quantification

Extracellular and intracellular lactate levels were quantitatively determined using a Lactic Acid assay kit (Nanjing Jiancheng Bioengineering Institute, Cat# A019-2-1) following the manufacturer’s instructions. Briefly, for extracellular lactate assessment, cell culture supernatants were collected from cells cultured at 1 × 10^6^ cells/well for 24 hours. For intracellular lactate analysis, cells were harvested, washed twice with PBS, and lysed in 200 µL of the provided lysis buffer. The lysates were centrifuged at 12,000 g for 10 minutes to obtain the supernatant. Both sample types were incubated with the reaction mixture containing lactate dehydrogenase and the chromogenic substrate for 30 minutes at 37°C. The optical density (OD) was measured at 450 nm using a microplate reader (Synergy HTX, BioTek). Relative lactate levels were normalized to the total protein concentration of each sample, which was determined using a BCA Protein Assay Kit (Share-bio, Cat# SB-WB013).

### Nuclear and Cytoplasmic Fractionation

Subcellular fractionation was performed to assess the nuclear translocation of proteins. Cells were washed with cold PBS and resuspended in 200 µL ice-cold Cytoplasmic Extraction Buffer (CEB) consisting of PBS supplemented with 0.1% NP-40, a Protease Inhibitor Cocktail (Share-bio, Cat# SB-WB026), and Phosphatase Inhibitors. The suspension was gently homogenized by pipetting up and down 15 times and incubated on ice for 5 minutes. Nuclei were pelleted by centrifugation at 12,000 g for 30 seconds at 4°C, and the supernatant was collected as the cytoplasmic fraction. The nuclear pellet was washed twice with 500 µL cold CEB to minimize cytoplasmic contamination and subsequently solubilized in 50 µL Nuclear Extraction Buffer (NEB) supplemented with 1% SDS and sonicated briefly on ice (3 cycles of 10s on/10s off). Fraction purity was rigorously validated by Western blot analysis using β-actin and Histone H3 as specific markers for the cytoplasmic and nuclear fractions, respectively

### Immunoprecipitation (Co-IP) and Western Blotting

For protein extraction, cells were lysed in RIPA Lysis and Extraction Buffer (Thermo Fisher Scientific, Cat# 89901) supplemented with a Protease Inhibitor Cocktail (Share-bio, Cat# SB-WB026) and Phosphatase Inhibitors on ice for 30 minutes. Lysates were clarified by centrifugation at 14,000× g for 15 minutes at 4°C. For Co-IP assays, 500 µg of total protein lysate was incubated with 1-2 µg of primary antibody or control IgG overnight at 4°C with gentle rotation. Subsequently, 30 µL of Protein A/G Magnetic Beads (Share-bio, Cat# SB-PR001) were added and incubated for 4 hours at 4°C. The immunocomplexes were washed five times with cold lysis buffer and eluted by boiling in 1× SDS Loading Buffer (Yeasen, Cat# 20315ES20) for 10 minutes.

For Western blotting, 20-30 µg of protein extracts were separated by SDS-PAGE and transferred onto polyvinylidene fluoride (PVDF) membranes (0.45 µm; Millipore). Membranes were blocked with 5% Albumin Bovine V (BSA; Solarbio, Cat# A8020) or 5% non-fat milk in TBST for 1 hour at room temperature. Membranes were then probed with primary antibodies overnight at 4°C, followed by incubation with HRP-conjugated Goat Anti-Rabbit (Share-bio, Cat# SB-AB0101) or Goat Anti-Mouse (Share-bio, Cat# SB-AB0102) secondary antibodies for 1 hour at room temperature. Protein bands were visualized using an enhanced chemiluminescence (ECL) system (ChemiDoc MP, Bio-rad). Band intensity was quantified using ImageJ software. A detailed list of primary antibodies and their dilution ratios is provided in Supplementary Table S1.

### Animal Model Studies

All animal procedures were conducted in strict accordance with the Guide for the Care and Use of Laboratory Animals and were approved by the Institutional Animal Care and Use Committee of Ren Ji Hospital (Approval No. 20250233). Male C57BL/6J mice (6-8 weeks old) were purchased from Shanghai Model Organisms Center, Inc. and housed under specific pathogen-free (SPF) conditions with a 12-hour light/dark cycle and free access to food and water.

For the generation of the orthotopic PDAC model, mice were anesthetized with isoflurane. A subcostal incision was made to expose the pancreas, and 2 × 10⁵ KPC1199 cells (or *Ldha*-knockdown variants as indicated) suspended in 20 µL of PBS were injected into the pancreatic tail. The incision was closed with sutures. Tumor progression was monitored weekly via bioluminescence imaging following intraperitoneal injection of D-Luciferin (Share-bio, Cat# SB-D1009, 150 mg/kg) using an IVIS Spectrum Imaging System (PerkinElmer).

For dietary lactate modulation experiments, mice were randomized into groups 7 days post-tumor implantation and administered 200 mM Sodium L-lactate (L-NaLac), Sodium D-lactate (D-NaLac), or Sodium Acetate (NaAce) dissolved in drinking water *ad libitum*. The pH of the drnking water was adjusted to 7.0, and solutions were refreshed every 2 days.

For the macrophage replacement model, endogenous macrophages were first depleted using liposomal clodronate. Mice received an initial intravenous injection of 200 µL Clodronate liposomes (Yeasen, Cat# 40337ES) or control PBS liposomes, followed by maintenance doses of 100 µL twice weekly via tail vein injection. Following the initial depletion, 2 × 10^6^ RAW264.7 cells (transduced with shNC, shBcl3, shBcl3+WT, or shBcl3+K21R) were adoptively transferred into the L-NaLac-treated tumor-bearing mice via tail vein injection every 3 days. Mice were euthanized at the experimental endpoint, and tumors were harvested for weight measurement and histological analysis.

### Real-Time Quantitative PCR (RT-qPCR)

Total RNA was extracted from cultured cells or homogenized tumor tissues using RNAiso Plus (Takara Bio Inc., Cat# 9109) following the manufacturer’s instructions. RNA concentration and purity were assessed using a NanoDrop spectrophotometer (Thermo Fisher Scientific). Reverse transcription was performed using 500 ng of total RNA with PrimeScript™ RT Master Mix (Takara Bio Inc., Cat# RR036A) in a 10 µL reaction volume. Quantitative PCR was carried out on an Applied Biosystems 7500 Real-Time PCR System using 2× Universal SYBR Green qPCR Premix (Share-bio, Cat# SB-Q204). The thermal cycling conditions included an initial denaturation step at 95°C for 30 seconds, followed by 40 amplification cycles of denaturation at 95°C for 5 seconds and annealing/extension at 60°C for 34 seconds. Relative gene expression was calculated using the 2^-ΔΔCt^ method, with *GAPDH* or *Actb* serving as the internal normalization control. The specific primer sequences for human and murine genes (*IL6*, *IL12B*, *ARG1*, *MRC2*, etc.) are listed in Supplementary Table S2.

### RNA Sequencing

Total RNA was extracted from the tissue samples using TRIzol reagent (Invitrogen, CA, USA) strictly following the manufacturer’s instructions. RNA purity and concentration were evaluated using a NanoDrop 2000 spectrophotometer (Thermo Scientific, USA), while RNA integrity was assessed using an Agilent 2100 Bioanalyzer (Agilent Technologies, Santa Clara, CA, USA). Only high-quality RNA samples (OD260/280 = 1.8-2.2, OD260/230 ≥ 2.0, RIN ≥ 7.0) were selected to construct the sequencing library. The cDNA libraries were prepared using the VAHTS Universal V6 RNA-seq Library Prep Kit according to the manufacturer’s protocol, where poly(A) mRNA was enriched using oligo(dT) magnetic beads, fragmented, and subjected to cDNA synthesis, end-repair, A-tailing, and adapter ligation. The transcriptome sequencing was conducted by OE Biotech Co., Ltd. (Shanghai, China) on an Illumina Novaseq 6000 platform, generating 150 bp paired-end reads. For data analysis, raw reads were processed using fastp software to remove low-quality reads and adapters. The obtained clean reads were mapped to the reference genome using HISAT2, and gene expression levels were calculated based on FPKM (Fragments Per Kilobase of transcript per Million mapped reads) with read counts obtained by HTSeq-count. Differential expression analysis was subsequently performed using DESeq2, with differentially expressed genes (DEGs) defined by a Q-value < 0.05 and log_2_(FoldChange) > 1.

### Untargeted Metabolomics Profiling

Metabolites were extracted from tissue samples using a cold methanol/water mixture, followed by ultrasonication and centrifugation at 12,000 rpm for 15 min at 4°C. The supernatant was evaporated to dryness under a gentle stream of nitrogen gas. To facilitate gas chromatography-mass spectrometry (GC-MS) analysis, samples underwent a two-step derivatization process: methoximation with methoxyamine hydrochloride in pyridine at 30°C for 90 min, followed by silylation with N-Methyl-N-(trimethylsilyl) trifluoroacetamide (MSTFA) at 37°C for 30 min. Analysis was performed on an Agilent GC system coupled to a mass spectrometer equipped with a DB-5MS capillary column. Helium was used as the carrier gas at a constant flow rate of 1.0 mL/min. The mass spectrometer was operated in electron impact (EI) ionization mode at 70 eV, scanning the mass range of m/z 50-600. Metabolites were identified by comparing mass spectra and retention times against the NIST reference library. Data normalization and pathway enrichment analyses were conducted using the MetaboAnalyst 6.0 platform.

### Single-Cell RNA-seq (scRNA-seq) Data Integration and Analysis

Publicly available PDAC scRNA-seq datasets were processed using the Seurat R package (v4.0) with standard quality control and normalization procedures as described above. To systematically evaluate the lactate metabolic potential across different cell phenotypes, a specific lactate transport gene signature was defined. This composite signature comprised five key transporter genes: *SLC16A3* (MCT4), *SLC16A1* (MCT1), *SLC16A7* (MCT2), *SLC5A8* (SMCT1), and *SLC5A12* (SMCT2). A module score was calculated for each individual cell based on the aggregate expression of this gene set using the AddModuleScore function in Seurat, which subtracts the aggregated expression of control feature sets to reduce noise. The resulting Lactate Transporter Scores were compared across annotated cell clusters within the tumor microenvironment. Cell populations containing fewer than 50 cells were excluded to ensure analytical robustness. Differences in signature scores between tumor and normal conditions were assessed and visualized. Data distributions are displayed using box plots on a log10-transformed y-axis to facilitate visualization of expression patterns across cell types with widely varying values.

To identify specific transporters mediating lactate uptake, we performed differential expression analysis between tumor-associated macrophages (Tumor group) and normal macrophages (Normal/Adjacent group). The Log2 Fold Change (Log_2_FC) for each gene was calculated as the log_2_-transformed ratio of the mean expression in the tumor group to that in the normal group (with a pseudocount added to avoid division by zero). Statistical significance was assessed using the non-parametric Wilcoxon rank-sum test. Results were visualized using lollipop charts to display Fold Changes and box plots to show expression distributions.

### Global Proteomic and L-Lactylome Profiling

For integrated proteomic analysis, proteins were extracted using a robust lysis buffer containing 8M Urea, 1% NP-40, and a cocktail of protease and deacetylase inhibitors. Lysates were sonicated, centrifuged, and quantified via the BCA assay. For global proteomics, proteins were reduced with DTT, alkylated with iodoacetamide, and digested with trypsin overnight. For L-lactylome profiling, tryptic peptides were specifically enriched using pan-anti-L-Lactyl Lysine antibody-conjugated agarose beads (PTM Biolabs), followed by rigorous washing and elution. Peptides were desalted using C18 ZipTips and subjected to high-resolution liquid chromatography-tandem mass spectrometry (LC-MS/MS) analysis. Experiments were performed on an Orbitrap Astral mass spectrometer coupled to a Vanquish Neo UHPLC system (Thermo Fisher Scientific). Data-independent acquisition (DIA) was performed with a scan range of m/z 350-1500, MS1 resolution of 240,000, and Higher-energy Collisional Dissociation (HCD) fragmentation energy set at 25 eV. Raw data were processed using DIA-NN (v1.8.1) or Spectronaut software. Protein identification and modification sites were filtered with a false discovery rate (FDR) of <1%. Bioinformatics analysis, including GO enrichment, KEGG pathway analysis, and motif analysis of lactylated sites, was performed to elucidate the functional networks regulated by lactate-dependent modifications.

### Bioinformatics Analysis

The Lactylation-Driven Macrophage Signature (LDMS) was established by identifying differentially expressed genes significantly downregulated in PDAC-CM-treated macrophages following p300 inhibition with C646. For clinical validation, transcriptome profiles (HTSeq-FPKM) and corresponding clinicopathological data were retrieved from the TCGA-PAAD cohort. Patients were stratified based on LDMS expression levels, and survival outcomes were assessed using Kaplan-Meier estimators and the Fleming-Harrington test. The tumor microenvironment landscape was characterized using the ESTIMATE algorithm to calculate Immune and Stromal Scores, while the relative abundance of infiltrating immune cell populations, including M2 macrophages, was deconvoluted via the CIBERSORT algorithm. Functional pathway activities, specifically regarding immune checkpoints, regulatory T cell signatures, and T cell co-inhibition, were quantified using Gene Set Variation Analysis (GSVA). Prognostic independence of the BKMS in the TMA cohort was evaluated using univariate and multivariate Cox proportional hazards regression models to calculate hazard ratios (HR) and 95% confidence intervals (CI). All statistical computations were performed using R software, with significance defined as P < 0.05.

### Protein Structure Prediction and Visualization

The amino acid sequences of the human P300 HAT domain (Uniprot ID: Q09472) and full-length BCL3 (Uniprot ID: P20749) were retrieved from the UniProt database. The three-dimensional structure of the P300-BCL3 complex was predicted using AlphaFold-Multimer via the AlphaFold Server (Google DeepMind). The prediction run generated five models, and the model with the highest confidence score (ipTM + pTM) was selected for further analysis. The predicted structure was visualized and rendered using PyMOL Molecular Graphics System (Version 2.x, Schrödinger, LLC). Intermolecular interactions, including hydrogen bonds and atomic distances between the BCL3 K21 residue and P300 catalytic residues, were analyzed using the “Measurement” wizard in PyMOL.

### Statistical Analysis

Statistical analyses were performed using GraphPad Prism software (GraphPad Software, Version 8.0). All in vitro experiments were conducted with at least three biologically independent replicates (n ≥ 3), and data are presented as the mean ± standard error of the mean (SEM). For comparisons between two independent groups, statistical significance was determined using unpaired, two-tailed Student’s *t*-tests. For comparisons involving more than two groups, one-way analysis of variance (ANOVA) was employed, followed by Tukey’s post-hoc test for multiple comparisons. For quantitative analysis of clinical specimens, the correlation between the BCL3-Kla Macrophage Score (BKMS) and other clinicopathological parameters was assessed using Pearson’s or Spearman’s correlation analysis based on data distribution. A *P*-value of < 0.05 was considered statistically significant.

## Result

### Integrated metabolomics and scRNA-seq identify TAMs as major lactate consumers in PDAC

To characterize the metabolic reprogramming landscape in PDAC, untargeted metabolomics profiling was conducted on a cohort of 20 paired tumor and adjacent non-tumor tissues (Figure 1A). Principal Component Analysis (PCA) and Orthogonal Partial Least Squares Discriminant Analysis (OPLS-DA) revealed distinct metabolic phenotypes between tumor and non-tumor tissues, indicating significant metabolic remodeling (Figure 1B). Volcano plots highlighted prominent metabolomic alterations (Figure 1C), while heatmaps visualized significantly downregulated or upregulated metabolites in tumors (Figure 1D). Functional enrichment analysis of the differential metabolites identified “Glycolysis/Gluconeogenesis” and “SLC-mediated transmembrane transport” as top enriched pathways (Figure 1E). Notably, while upstream glycolytic intermediates (e.g., Glucose-6-Phosphate) accumulated robustly and lipid metabolism was suppressed, lactate levels did not show a commensurate increase in tumor tissues (Figures 1F and 1G). Given the enrichment of transport pathways, we hypothesized that this apparent lactate deficit stems not from reduced production, but from rapid turnover and consumption by specific cellular components within the TME.

**Figure 1.**
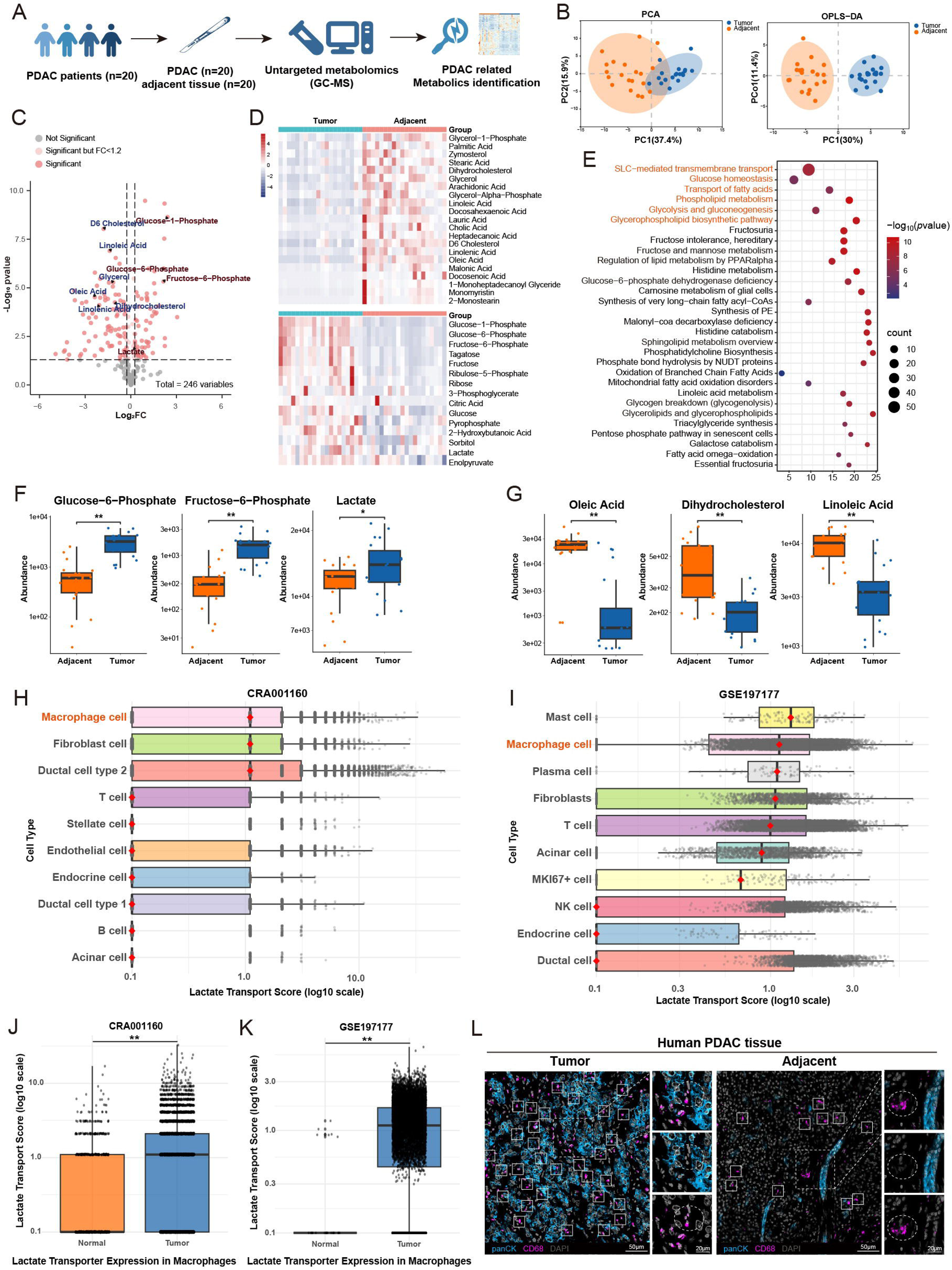
Distinct metabolic landscapes and cell-type specific lactate transporter patterns in PDAC. **(A)** Schematic workflow of untargeted metabolomics profiling performed on paired PDAC tumor (T) and adjacent normal (N) tissues (n = 20). **(B)** Multivariate analysis using Principal Component Analysis (PCA) and Orthogonal Partial Least Squares Discriminant Analysis (OPLS-DA) score plots to compare metabolic profiles between T and N groups. **(C-D)** Identification of differential metabolites. **(C)** Volcano plot showing the distribution of metabolites based on fold change and *P*-value (red dots indicate significant alterations, *P* < 0.05). **(D)** Heatmap visualizing the hierarchical clustering of significantly downregulated (upper) and upregulated (lower) metabolites (color scale: z-score of relative abundance). **(E)** Pathway enrichment analysis of differential metabolites performed using MetaboAnalyst 6.0. **(F-G)** Relative abundance of representative metabolites involved in glycolysis **(F)** and lipid metabolism **(G)** (mean ± SEM, paired two-tailed *t* test, n = 20 pairs). **(H-I)** Distribution of the composite lactate transporter expression signature across major cell populations in the PDAC microenvironment from the CRA001160 **(H)** and GSE197177 **(I)** datasets. Each box plot represents the signature score per cell type; the red diamond denotes the median value. Data are presented on a log10 scale. **(J-K)** Comparative analysis of the lactate transporter signature within macrophage subsets between matched normal and tumor tissues from the CRA001160 **(J)** and GSE197177 **(K)** datasets. **(L)** Representative multiplex immunofluorescence (mIF) images displaying the spatial distribution of macrophages (CD68^+^) and tumor cells (pan-CK^+^) in human PDAC tissues. \**P* < 0.05; \*\**P* < 0.01.

To identify the cellular sink for lactate within the TME of PDAC, we integrated public scRNA-seq datasets (CRA001160 and GSE197177) and constructed a “Lactate Transporter Score” based on the aggregate expression of five key transporters (*SLC16A3*, *SLC16A1*, *SLC16A7*, *SLC5A8*, and *SLC5A12*). Cross-cell-type comparison identified macrophages as the predominant population expressing high-affinity lactate transporters, significantly outranking epithelial or other stromal cells (Figures 1H and 1I). Moreover, lactate transporter scores were markedly elevated in TAMs compared to their normal tissue counterparts (Figures 1J and 1K). mIF analysis of human PDAC tissues revealed extensive macrophage infiltration proximal to tumor cells (Figure 1L). Collectively, these multi-omics results imply that TAMs could function as a key lactate consumer within the TME, providing a plausible explanation for the observed disconnect between glycolytic activity and lactate accumulation.

### PDAC-derived lactate drives M2-like macrophage polarization *in vitro* and *in vivo*

To recapitulate tumor-immune interactions *in vitro*, we exposed PBMC-derived macrophages to PDAC cells (co-culture) or PDAC-conditioned medium (CM) (Figure S1A). Exposure to PDAC-derived factors triggered a morphological transition toward an elongated, spindle-like architecture in PBMC-derived macrophages (Figures S1B and S1C). To determine whether this morphological transformation corresponded to a specific functional polarization state, we analyzed canonical macrophage markers. qPCR analysis revealed that these macrophages acquired a distinct tumor-promoting phenotype, characterized by the robust upregulation of M2 markers (*ARG1*, *MRC2*, and *IL10*) (Figure S1D). Consistent with *in vitro* data, quantitative assessment of human PDAC tissues showed a specific enrichment of macrophages co-expressing CD16, ARG1, and CD86 (Figure 2A). Importantly, the phenotypic remodeling induced by PDAC coculture or CM was conserved across human (THP-1) and murine (RAW264.7) macrophages (Figures 2B, S1E and S1G) and extended to their migratory potential (Figures S1F and S1H).

**Figure 2.**
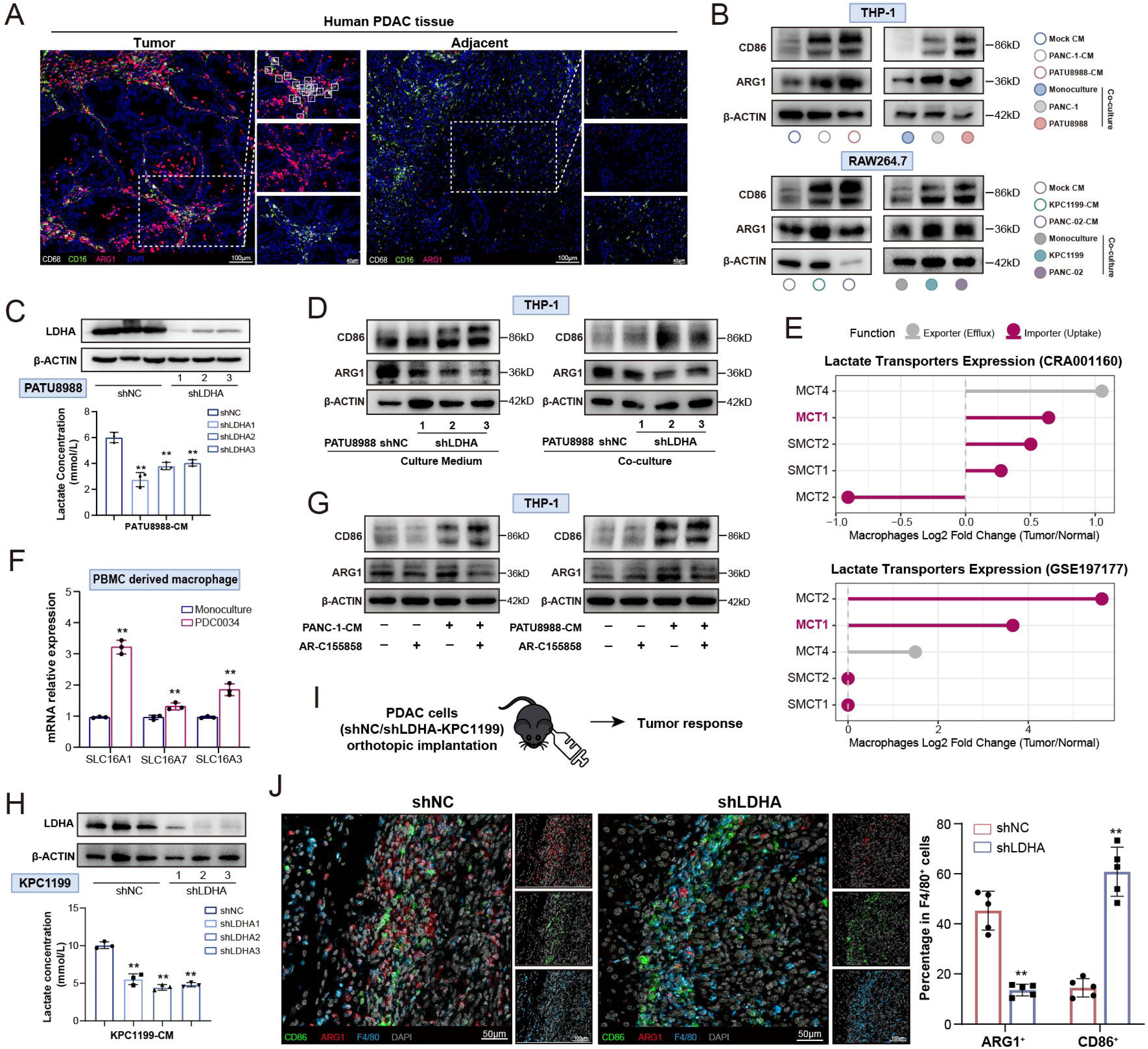
PDAC-derived lactate drives M2-like polarization of macrophages via MCT1-mediated transport. **(A)** Multiplex IF visualization and identification of CD16^+^ARG1^+^CD86^+^ macrophage subpopulations in paired PDAC and adjacent normal tissues. **(B)** Immunoblot analysis of ARG1 and CD86 expression in human (THP-1) and murine (RAW264.7) macrophages under co-culture conditions or CM stimulation. **(C)** Verification of *LDHA* knockout efficiency and measurement of lactate concentrations in supernatants of PATU8988 cells (mean ± SEM, unpaired two-tailed *t* test, n = 3 per group). **(D)** Immunoblot analysis of polarization markers in THP-1 macrophages co-cultured with control or *LDHA*-deficient tumor cells. **(E)** Differential expression of lactate transporters in macrophages (CRA001160 and GSE197177). Data are presented as Log2 fold change (Tumor vs. Normal). Colors indicate transporter function (Red: Importer; Gray: Exporter). **(F)** Relative mRNA expression of *SLC16A1* (MCT1), *SLC16A7* (MCT2), and *SLC16A3* (MCT4) in primary macrophages following co-culture with PDAC cells (mean ± SEM, unpaired two-tailed *t* test, n = 3 per group). **(G)** Assessment of polarization markers at protein levels in THP-1 cells stimulated with PDAC-CM ± AR-C155858. **(H)** Validation of *Ldha*-KD efficiency and lactate concentration measurement in KPC1199 cells supernatants (mean ± SEM, unpaired two-tailed *t* test, n = 3 per group). **(I)** Schematic illustration of the orthotopic mouse model utilizing *Ldha*-KD KPC1199 cells. **(J)** Representative mIF staining of orthotopic tumor sections quantifying the ratio of ARG1^+^F4/80^+^ versus CD86^+^F4/80^+^ subpopulations (mean ± SEM, unpaired two-tailed *t* test, n = 5 per group). \**P* < 0.05; \*\**P* < 0.01.

Given the hyper-glycolytic phenotype of PDAC, we hypothesized that tumor-derived lactate acts as a key metabolic driver. Metabolic quantification confirmed elevated lactate concentrations in PDAC culture supernatants (Figure S2A). To specifically block lactate production, we targeted Lactate Dehydrogenase A (LDHA), the terminal glycolytic enzyme converting pyruvate to lactate.^22^ LDHA depletion significantly reduced lactate secretion in PDAC cells (Figure 2C) and abrogated the induction of macrophage ARG1 expression while enhancing CD86 levels (Figures 2D and S2B). Notably, LDHA deficiency did not alter macrophage migration (Figure S2C), suggesting that while lactate drives polarization, other factors in the CM may mediate recruitment. Pharmacological LDH inhibition (PX-478 or GSK-2837808A) similarly reduced supernatant lactate (Figure S2D).

To identify the specific transporter mediating lactate entry into macrophages, we interrogated scRNA-seq datasets (CRA001160 and GSE197177). Based on the premise that macrophages may be major consumers of tumor-derived lactate, we prioritized high-affinity transporters responsible for lactate uptake. While the efflux transporter *SLC16A3* (MCT4) was upregulated, reflecting the glycolytic nature of TAMs, *SLC16A1* (MCT1) emerged as the predominant high-affinity importer in TAMs compared to normal counterparts across both databases (Figure 2E). Consistent with these bioinformatic findings, *in vitro* co-culture with PDAC cells robustly upregulated *SLC16A1* (MCT1) expression in PBMC-derived macrophages (Figure 2F). Subsequent blockade of MCT1/2-mediated uptake (AR-C155858) prevented PDAC-induced morphological transitions (Figure S2E), suppressed intracellular lactate accumulation (Figures S2F and S2H), and impaired the acquisition of the M2-like phenotype (Figures 2G, S2G and S2I). Conversely, inhibition of lactate efflux via 7ACC1 had opposing effects on intracellular lactate accumulation and M2 polarization (Figures S2J-M).

To validate these findings *in vivo*, we established an orthotopic PDAC model by implanting *Ldha*-knockdown (*Ldha*-KD) KPC1199 murine PDAC cells into the pancreas of C57BL/6J mice (Figures 2H and 2I). Bioluminescence imaging and tumor weight analysis confirmed that *Ldha* deficiency significantly attenuated tumor burden (Figure S2N), a result attributable to both restricted tumor glycolysis and TME dysregulation.^23^ Notably, immune profiling revealed that restriction of lactate production drove a phenotypic shift from a pro-tumor ARG1^+^ state to a pro-inflammatory CD86^+^ state (Figure 2J). Supernatants from *Ldha*-KD KPC1199 clones failed to induce ARG1 expression in RAW264.7 macrophages (Figure S2O), and consequently lacked the capacity to promote KPC1199 proliferation as evidenced by EdU assays (Figure S2P). Collectively, these findings identify PDAC-secreted lactate as a pivotal driver of macrophage reprogramming toward a tumor-promoting phenotype.

### Lactate reprograms macrophages toward a tumor-promoting phenotype to facilitate PDAC progression

Supplementation of PDAC-CM with exogenous lactate further potentiated intracellular lactate accumulation and M2-associated gene expression in RAW264.7 macrophages (Figures S3A). However, unlike PDAC-CM, lactate alone failed to promote migration, indicating that lactate primarily drives macrophage reprogramming rather than chemotaxis (Figures S3B and S3C). Phenotypically, lactate induced a robust shift toward a tumor-promoting state in macrophages, marked by ARG1 upregulation and pro-inflammatory marker suppression (Figures 3A, 3B, S3D and S3E). Treatment with AR-C155858, an inhibitor for lactate uptake, effectively abrogated lactate-induced phenotypic reprogramming in both human (THP-1) and murine (RAW264.7) macrophages (Figures S3F and S3G). Functionally, CM from lactate-primed macrophages (Lac-PrM) significantly promoted malignant proliferation of both human (PANC-1, PATU8988) and murine (KPC1199, PANC-02) PDAC cells (Figure 3C-D). To validate above findings *in vivo*, Lac-PrM were adoptively transferred into mice bearing orthotopic KPC1199 tumors (Figure 3E). The adoptive transfer significantly accelerated tumor growth and proliferation, evidenced by bioluminescence imaging and increased tumor weights (Figure 3F). Histological analysis confirmed enhanced tumor cell proliferation (Ki67+) (Figure 3G) and an M2-like TAMs profile (increased ARG1^+^F4/80^+^ to CD86^+^F4/80^+^ ratio) (Figure 3H).

**Figure 3.**
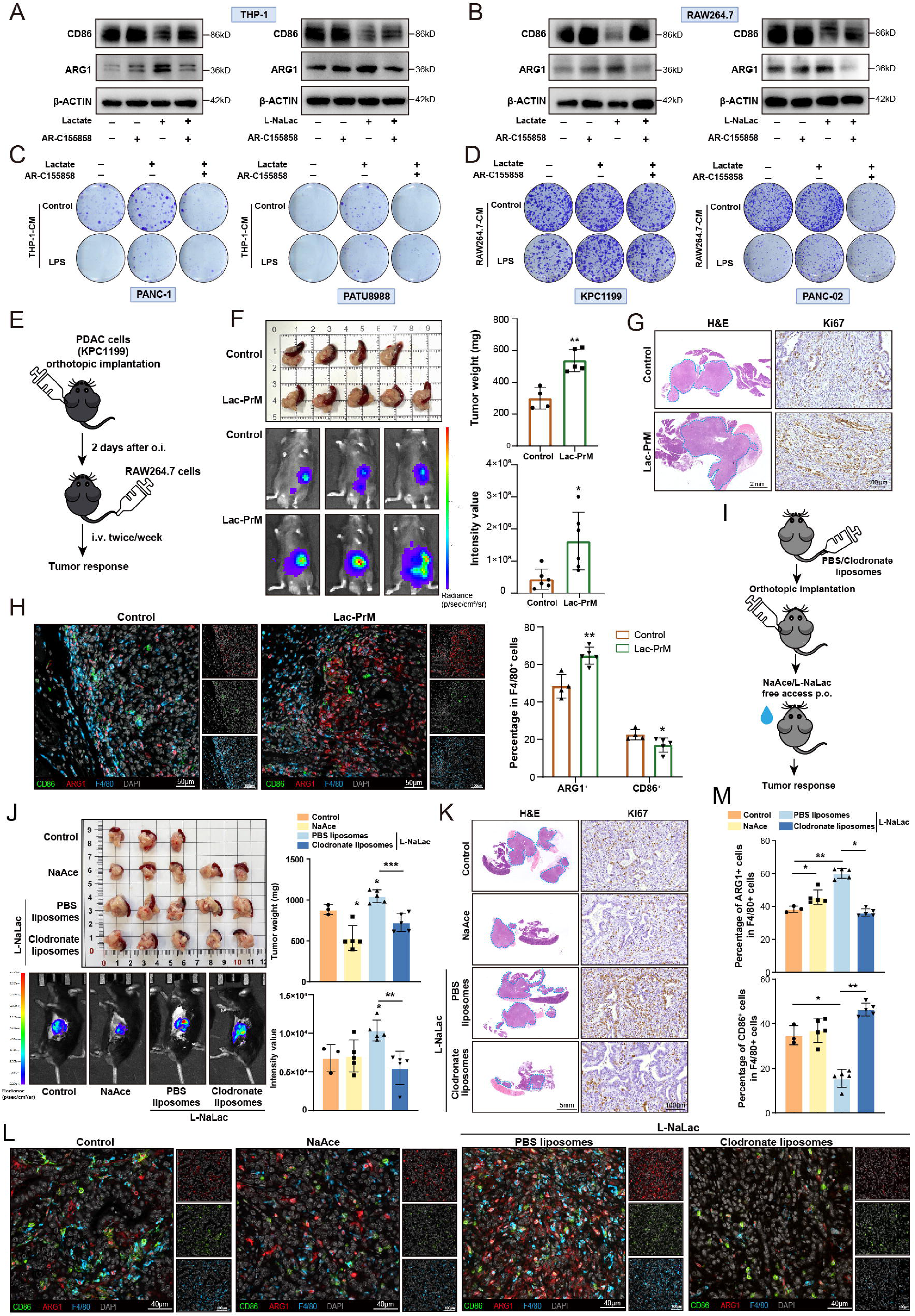
Exogenous lactate enforces a tumor-promoting macrophage phenotype to accelerate PDAC progression. (A-B) Immunoblot analysis of ARG1 and CD86 expression in THP-1 **(A)** and RAW264.7 **(B)** macrophages treated with lactate or L-sodium lactate in the presence or absence of AR-C155858. **(C-D)** Colony formation assays assessing the proliferation of human (PANC-1, PATU8988) **(C)** and murine (KPC1199, PANC-02) **(D)** PDAC cells cultured with CM derived from lactate- or L-sodium lactate-primed macrophages. **(E)** Schematic illustration of the orthotopic PDAC mouse model involving adoptive transfer of lactate-primed macrophages (Lac-PrM) via tail vein injection. **(F)** Quantification of tumor burden via bioluminescence imaging and terminal tumor weights. (mean ± SEM, unpaired two-tailed *t* test, n = 4 for Control, n = 5 for Lac-PrM). **(G)** Representative H&E staining and Ki67 IHC of orthotopic tumor tissue sections. **(H)** Representative mIF staining and quantification of ARG1^+^F4/80^+^ versus CD86^+^F4/80^+^ macrophage subpopulations (mean ± SEM, unpaired two-tailed *t* test). **(I)** Schematic illustration of the orthotopic PDAC model combined with pharmacological macrophage depletion. Treatment groups include vehicle, NaAce, or L-NaLac combined with either PBS liposomes (PBS-Lip) or clodronate liposomes (Clo-Lip). **(J)** Assessment of tumor burden via bioluminescence imaging and measurement of terminal tumor weights (mean ± SEM, one-way ANOVA with Tukey’s test, n = 3 for Control, n = 5 for NaAce, n = 5 for L-NaLac + PBS-Lip, n = 5 for L-NaLac + Clo-Lip). **(K)** Representative H&E staining and IHC for Ki67 in orthotopic PDAC tissue sections. **(L-M)** Representative mIF images **(L)** and quantitative analysis **(M)** of polarization status (ARG1^+^F4/80^+^ vs. CD86^+^F4/80^+^) within the TME (mean ± SEM, one-way ANOVA with Tukey’s test). \**P* < 0.05; \*\**P* < 0.01.

To determine whether these effects were driven by specific stereoselective signaling or by non-specific acidification/carbon source utilization, we utilized the stereoisomer D-sodium lactate (D-NaLac) and sodium acetate (NaAce) as controls.^24^ *In vitro*, neither induced significant migration or phenotypic polarization (Figures S3H-K). Crucially, direct administration of lactate to PDAC cells did not alter proliferation *in vitro* (Figure S3L and S3M). Consistently, oral administration of L-NaLac to orthotopic KPC1199 tumor-bearing mice significantly exacerbated tumor burden, whereas NaAce and D-NaLac had no observable effect (Figures S3N and S3O). Correspondingly, histological and mIF analyses revealed that L-NaLac specifically enhanced tumor cell proliferation (Ki67^+^), and promoted M2-like polarization (ARG1^+^), while NaAce and D-NaLac failed to trigger such phenotypes (Figures S3P-R). These data confirmed that tumor-promoting effects are mediated indirectly through macrophage reprogramming via an enantiomer-dependent, acidification-independent mechanism

To further determine the requirement for macrophages in lactate-driven tumor progression, we depleted macrophages using clodronate liposomes in the orthotopic PDAC model (Figure 3I). Consistent with *in vitro* findings, oral administration of L-NaLac significantly exacerbated tumor burden, whereas NaAce failed to induce such changes (Figure 3J). However, macrophage depletion effectively abrogated the tumor-promoting effects of exogenous L-NaLac (Figure 3J). Histological analysis (H&E, Ki67) confirmed that lactate-induced tumor proliferation is abolished in the absence of macrophages (Figure 3K). mIF analysis further demonstrated that lactate administration promotes a shift toward a M2-like phenotype (ARG1^+^F4/80^+^) compared to vehicle controls (Figures 3L and 3M). These observations indicate that TAMs are indispensable mediators of lactate-driven PDAC progression.

### Tumor-derived lactate drives macrophage reprogramming via protein L-lactylation

The stereoselective requirement for L-lactate implicated a specific signaling role rather than non-specific acidification. Therefore, we reasoned that protein L-lactylation (Kla) might serve as a molecular link between metabolic flux and transcriptional regulation.^12,25^ mIF analysis validated elevated Kla levels in macrophages within PDAC tissues compared to adjacent normal tissues (Figure 4A). Co-culture studies similarly demonstrated Kla upregulation in PDAC-exposed macrophages (Figure 4B). Notably, LDHA depletion abolished macrophage Kla induction in both *in vitro* co-cultures system and *in vivo* orthotopic tumors, whereas acetylation levels remained unaffected *in vitro* (Figures 4C and 4D). Pharmacological blockade of the transporter MCT1/2 (AR-C155858) suppressed Kla in macrophages (Figure 4E). In contrast, Lac-PrM maintained robustly elevated Kla levels following adoptive transfer into orthotopic tumors, indicating that Kla accumulation is strictly dependent on direct lactate uptake (Figure 4F).

**Figure 4.**
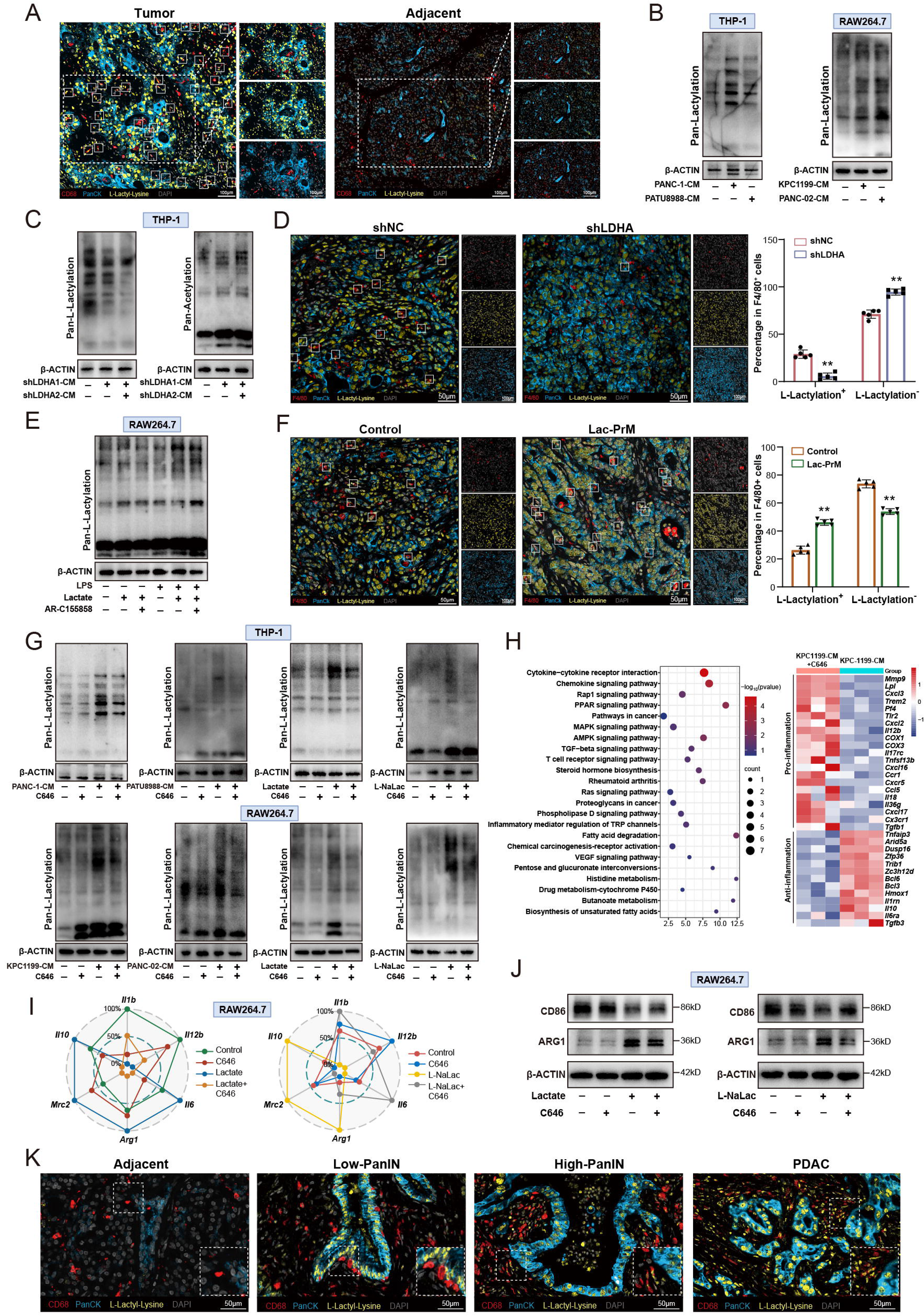
Protein L-lactylation couple metabolic status to macrophage phenotypic reprogramming. **(A)** Representative mIF visualizing the co-localization of Pan-Kla (L-lactylation) and CD68 (macrophage marker) in paired human PDAC and adjacent normal tissues. **(B)** Immunoblot analysis of global protein L-lactylation (Pan-Kla) levels in THP-1 and RAW264.7 macrophages co-cultured with the indicated PDAC cell lines. **(C)** Immunoblot analysis of Pan-Kla (Left) and Pan-Acetyl (Right) levels in THP-1 macrophages co-cultured with control or *LDHA*-deficient PATU8988 cells. **(D)** Analysis of macrophage L-lactylation levels in orthotopic tumors derived from control versus shLDHA cells. **(E)** Immunoblot analysis of Pan-Kla in RAW264.7 cells treated with AR-C155858. **(F)** Evaluation of macrophage L-lactylation status in orthotopic tumors following adoptive transfer of lactate-primed macrophages (Lac-PrM) (mean ± SEM, unpaired two-tailed *t* test, n = 5 per group). **(G)** Immunoblot analysis of Pan-Kla in THP-1 and RAW264.7 cells stimulated with PDAC-CM, lactate, or L-sodium lactate ± C646. **(H)** Functional enrichment analysis and heatmaps of immune-related gene expression derived from RNA-sequencing of RAW264.7 cells stimulated with KPC1199 CM ± C646. **(I-J)** Radar plots of immune-associated gene expression **(I)** and ARG1/CD86 levels **(J)** in RAW264.7 cells stimulated with lactate, or L-sodium lactate ± C646. **(K)** Representative mIF images illustrating macrophage L-lactylation dynamics across human PDAC progression (Normal, PanIN, and PDAC). \**P* < 0.05; \*\**P* < 0.01.

To distinguish the effects of extrinsic versus intrinsic lactate within the mixed M1/M2 polarization state of the TME, we compared exogenous lactate exposure with LPS stimulation, which induces glycolysis and M1 polarization.^6^ Despite enhancing endogenous lactate production, LPS failed to elevate Kla levels (Figure S4A), suggesting that the modification is primarily driven by uptake of exogenous, tumor-derived lactate rather than by intrinsic metabolic shifts.

To investigate a causal relationship between lactylation and phenotypic remodeling, we focused on p300, which has been identified as a key “writer” enzyme responsible for protein lactylation.^26,27^ Treatment with C646, a p300 inhibitor, not only effectively abrogated Kla induction triggered by PDAC-CM, exogenous lactate, or L-NaLac (Figure 4G) but also reversed the lactate-induced gene signature, abolishing ARG1 expression while restoring pro-inflammatory markers (Figures 4H, 4J and S4B). Functionally, preventing lactylation via C646 eliminated the capacity of Lac-PrM to support PDAC cell proliferation (Figures S4C-G). Spatiotemporal characterization further revealed that Kla levels increase progressively from precursor PanIN lesions to invasive PDAC, with specific enrichment in macrophages proximal to malignant PDAC cells (Figure 4K). Collectively, these data suggest that protein L-lactylation functions as a key molecular link, translating tumor-derived metabolic signals into pro-tumorigenic macrophage reprogramming.

### Integrated profiling identifies BCL3 L-lactylation as a driver of NF-κB-mediated macrophage reprogramming

To identify downstream effectors driving TAMs polarization, we performed an integrated analysis of quantitative global proteomics and L-lactylome profiling on KPC1199-CM-treated RAW264.7 macrophages (Figure 5A). Global proteomic analysis revealed differentially expressed proteins significantly enriched in inflammation-associated signaling including NF-κB and TNF pathways (Figure S5A). Analysis of the L-lactylome data revealed that differentially lactylated proteins were predominantly localized to the nucleus (Figure S5B) and included numerous transcription factors such as BCL3, Dnmt1, and Junb (Figures S5C). Notably, protein domain enrichment analysis identified a significant enrichment of the Ankyrin repeat domain (ARD) among differentially modified proteins (Figure S5D). Furthermore, KEGG pathway analysis confirmed that these lactylated proteins, particularly the nuclear fraction, were significantly enriched in inflammatory response and immune regulation pathways (Figures S5E and S5F).

**Figure 5.**
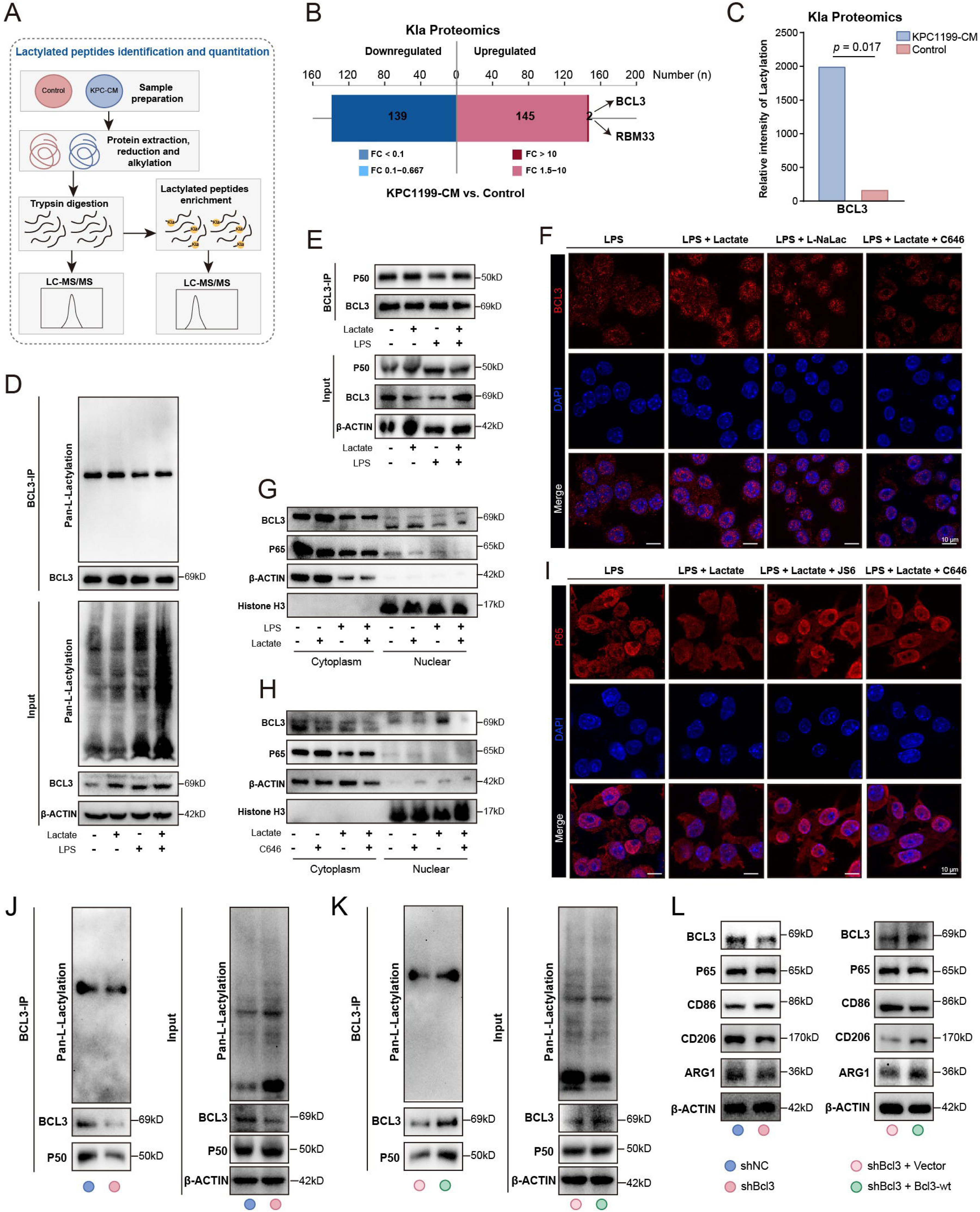
Integrated profiling identifies BCL3 L-lactylation as a driver of NF-κB-mediated macrophage reprogramming. **(A)** Schematic workflow for the integrated quantitative global proteomic and L-lactylation proteomic profiling of RAW264.7 macrophages stimulated with KPC1199-CM. **(B)** Distribution plot displaying differentially modified proteins ranked by enrichment fold change. **(C)** Density distribution analysis comparing BCL3 L-lactylation intensity between groups. **(D)** Co-IP assays assessing BCL3 L-lactylation levels (Pan-Kla) in RAW264.7 cells stimulated with lactate. **(E)** Co-IP analysis of the endogenous interaction between BCL3 and the NF-κB p50 subunit in RAW264.7 cells stimulated with lactate. **(F)** Representative ICC images visualizing BCL3 nuclear translocation in RAW264.7 cells treated with lactate ± C646. **(G-H)** Immunoblot analysis of BCL3 and NF-κB p65 in cytosolic versus nuclear fractions of RAW264.7 cells stimulated with lactate **(G)** or lactate ± C646 **(H)**. **(I)** Representative ICC images of NF-κB p65 localization in RAW264.7 cells treated with lactate ± C646 or JS6. **(J-K)** Co-IP and immunoblot analysis of pan-L-lactylation and p50 interaction in *Bcl3*-KD **(J)** and reconstituted **(K)** RAW264.7 cells under lactate stimulation. **(L)** Immunoblotting analysis of phenotypic markers (CD86, CD206, ARG1) in *Bcl3*-deficient and rescued RAW264.7 cells under lactate stimulation.

By applying a multi-dimensional filtering strategy that prioritized nuclear localization, high enrichment fold change, the presence of an ARD, and immune-related functional annotation, the atypical IκB family member BCL3 emerged as the top candidate (Figure 5B). Density distribution analysis further confirmed a marked increase in BCL3 L-lactylation intensity in the treated group (Figure 5C). We next investigated the molecular mechanisms underlying L-lactylation-mediated BCL3 regulation. Co-immunoprecipitation (Co-IP) assays confirmed that BCL3 L-lactylation was robustly induced by exogenous lactate but markedly suppressed by the MCT1/2 inhibitor AR-C155858 (Figures 5D and S6A), indicating that BCL3 modification is strictly dependent on MCT-mediated transport of exogenous lactate.

Given that BCL3 functions as an atypical IκB family member and transcriptional co-regulator, we examined its interaction with NF-κB subunits.^17,28^ Co-IP analysis demonstrated that lactate treatment markedly enhanced the endogenous interaction between BCL3 and the NF-κB p50 subunit (Figure 5E). Crucially, immunocytochemistry (ICC) and subcellular fractionation revealed that lactate exposure triggers BCL3 nuclear translocation, which coincides with a reduction in the nuclear abundance of the NF-κB p65 (RelA) subunit (Figures 5F and 5G). Pharmacological inhibition of lactylation via C646 abolished BCL3 nuclear entry and restored p65 nuclear accumulation (Figures 5F and 5H). Consistently, ICC analysis demonstrated that either preventing lactylation (C646) or disrupting the BCL3-NF-κB interaction (JS6) was sufficient to restore p65 nuclear localization (Figure 5I). Together, these findings support a mechanism wherein lactylated BCL3 competes for p50 binding, potentially sequestering the p50 subunit and thereby limiting the nuclear translocation and transcriptional activity of the canonical p65/p50 heterodimer.

Furthermore, JS6-mediated disruption of the BCL3-NF-κB complex in PDAC-CM or lactate-stimulated macrophages abolished M2 (ARG1) induction while restoring M1 (CD86) expression (Figure S6B). Consistent with these protein-level changes, qPCR analysis confirmed that disrupting this complex reverses the pro-tumor gene signature to a pro-inflammatory profile (Figure S6C). Finally, to verify the requirement for BCL3 in lactate-driven reprogramming, we utilized *Bcl3*-deficient cells. Lactate stimulation failed to induce BCL3 lactylation or p50 interaction in *Bcl3*-KD cells, but BCL3 re-expression fully rescued these defects (Figures 5J and 5K). Functionally, Bcl3 silencing in RAW264.7 macrophages impaired lactate-driven M2-like polarization, whereas re-expression restored the polarization state (Figure 5L), indicating BCL3 as an essential mediator of lactate-induced macrophage reprogramming.

### Site-specific L-lactylation at K21 governs BCL3-NF-κB interaction and macrophage pro-tumor reprogramming

To map the precise amino acid residue responsible for BCL3 L-lactylation, we employed high-resolution LC-MS/MS analysis, which identified Lysine 21 (K21) as the primary modification site (Figure 6A). To explore the molecular basis of this modification, we utilized AlphaFold-Multimer to model the interaction interface between the P300 catalytic HAT domain and BCL3. The predicted model suggested a binding pose wherein the N-terminal region of BCL3 is accommodated within the substrate-binding cleft of P300 (Figure S5G, left). Notably, visualization of the catalytic center indicated that the BCL3 K21 residue extends into the enzymatic pocket, potentially forming a hydrogen bond with the Aspartate 158 (D158) residue of P300 at a distance of 2.6 Å (Figure S5G, right). These structural data provide a potential mechanistic basis for P300-mediated K21 lactylation. Of note, structural modeling places the K21 residue within the N-terminal regulatory domain, which harbors the essential nuclear localization sequence (NLS) (Figure 6B),^29,30^ suggesting that K21 modification may influence BCL3 nuclear entry. The functional relevance of K21 was supported by evolutionary analysis, which demonstrated that the target lysine and its flanking sequences are highly conserved across species (Figures S5H and S5I).

**Figure 6.**
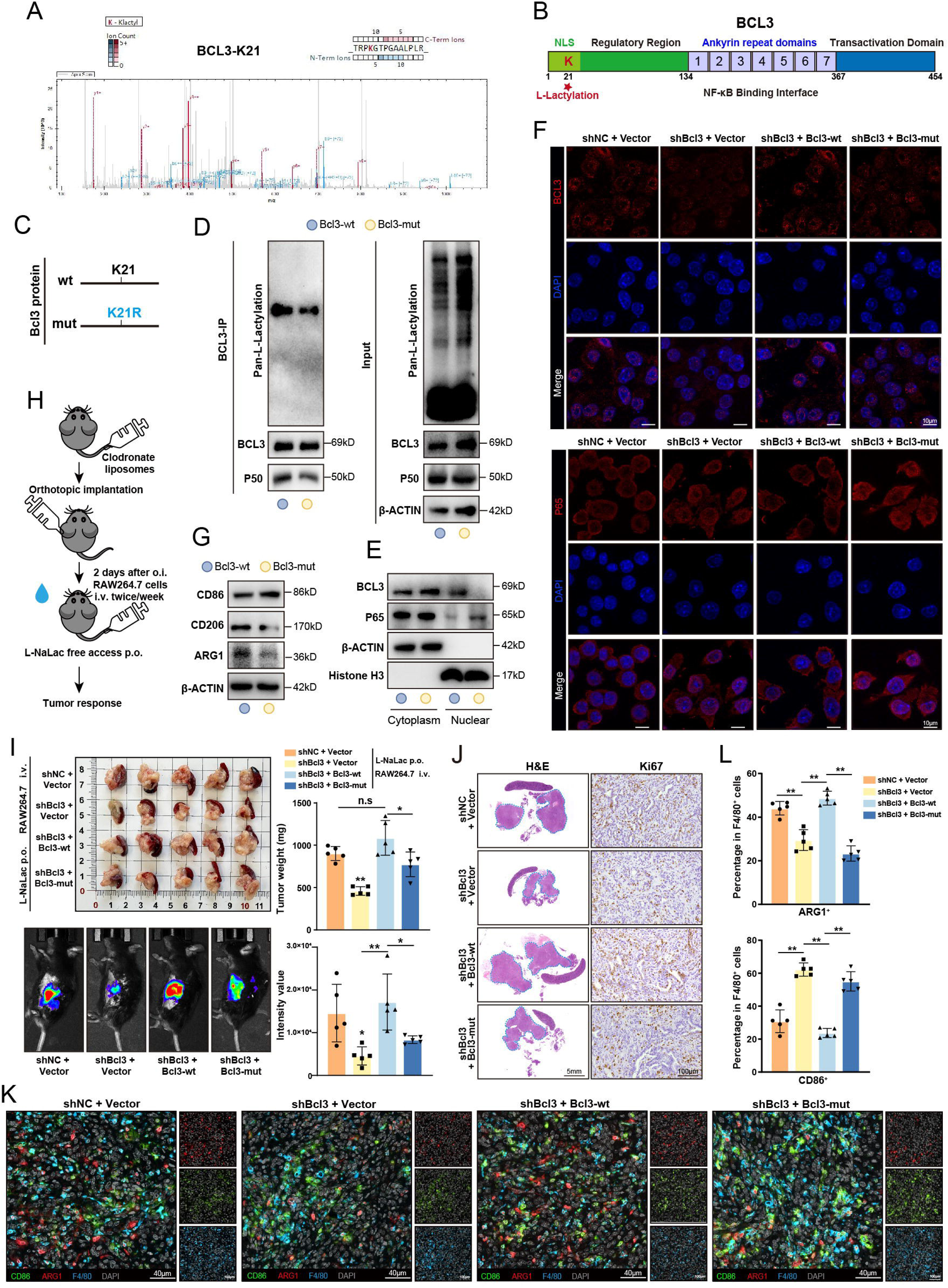
Site-specific L-lactylation at K21 governs BCL3-NF-κB interaction and macrophage pro-tumor reprogramming. **(A)** Annotated MS/MS spectrum identifying Lysine 21 (K21) as the primary L-lactylation site of BCL3. **(B)** Structural modeling of BCL3 interacting with the NF-κB dimer, highlighting the K21 residue within the NLS domain. **(C)** Schematic illustration of the site-directed mutagenesis strategy for generating the lactylation-deficient BCL3 K21R mutant. **(D)** Co-IP analysis of BCL3 L-lactylation and p50 interaction in WT versus K21R mutant RAW264.7 cells. **(E)** Immunoblot analysis of BCL3 and p65 in cytosolic and nuclear fractions in WT and K21R mutant RAW264.7 cells. **(F)** Representative ICC staining of BCL3 and p65 localization in WT and K21R mutant RAW264.7 cells. **(G)** Immunoblotting of polarization markers (CD86, CD206, ARG1) in RAW264.7 cells expressing WT or K21R mutant BCL3 treated with lactate. **(H)** Schematic workflow for the “macrophage replacement” model. Endogenous macrophages were depleted via Clo-Lip followed by adoptive transfer of RAW264.7 cells (*Bcl3*-KD, WT-Rescue, K21R-Rescue) into L-NaLac-treated PDAC mice via tail vein injection. **(I)** Quantification of tumor burden via bioluminescence imaging and terminal tumor weights (mean ± SEM, one-way ANOVA with Tukey’s test, n = 5). **(J)** Representative H&E staining and IHC for Ki67 in orthotopic PDAC tissues. **(K-L)** Representative mIF images **(K)** and quantification **(L)** of macrophage polarization phenotypes (ARG1^+^F4/80^+^ vs. CD86^+^F4/80^+^) (mean ± SEM, one-way ANOVA with Tukey’s test). \**P* < 0.05; \*\**P* < 0.01.

To interrogate the specific function of the K21 site, *Bcl3*-KD macrophages were reconstituted with wild-type (WT) or lactylation-deficient K21R mutant Bcl3 (Figures 6C and S6D). Mechanistically, the K21R mutation markedly attenuated the lactate-induced BCL3-p50 interaction (Figure 6D). Nuclear/cytosolic fractionation and ICC analyses revealed that unlike the WT protein, the K21R mutant is predominantly retained in the cytoplasm, failing to sequester p50 and thereby permitting nuclear translocation of the pro-inflammatory p65 subunit (Figures 6E and 6F). Functionally, macrophages expressing the K21R mutant exhibited blunted M2 marker upregulation (ARG1, CD206) and failed to suppress pro-inflammatory genes under lactate stimulation (Figure 6G and S6E). Moreover, neither *Bcl3*-KD nor K21R-expressing macrophages supported PDAC cell proliferation (KPC1199 and PANC-02), mirroring the growth-suppressive effects of the BCL3 inhibitor JS6, as demonstrated by colony formation and EdU assays (Figures S6F and S6G). Together, these findings imply that L-lactylation at K21 governs BCL3 nuclear transport, p50/p65 complex regulation, and subsequent tumor-promoting capabilities.

To verify the functional necessity of the BCL3 K21 site *in vivo*, we utilized a macrophage replacement strategy wherein endogenous macrophages were depleted and replaced via adoptive transfer of *Bcl3*-KD, WT-rescue, or K21R-rescue RAW264.7 cells into L-NaLac-fed PDAC mice (Figure 6H). Consistent with *in vitro* findings, mice receiving Bcl3-deficient macrophages exhibited markedly reduced tumor burden compared to controls (Figure 6I). Rescue experiments confirmed that while re-introduction of WT BCL3 fully restored macrophage-driven tumor growth, the lactylation-deficient K21R mutant failed to recapitulate the tumor-promoting capacity, resulting in tumor burdens comparable to the *Bcl3*-KD group (Figures 6I and 6J). Crucially, quantitative mIF analysis demonstrated that while WT BCL3 enabled lactate-induced polarization toward an ARG1^+^ M2-like state, macrophages expressing the K21R mutant remained resistant to the reprogramming, retaining a predominant CD86^+^ phenotype despite systemic lactate supplementation (Figures 6K and 6L). Collectively, the *in vivo* data demonstrate that BCL3 L-lactylation at K21 is a critical determinant for macrophage reprogramming and the subsequent promotion of PDAC progression.

### Lactylation-driven macrophage reprogramming correlates with poor prognosis and an immunosuppressive TME in PDAC

To evaluate the clinical relevance of lactate-driven macrophage reprogramming, we derived a “Lactylation-Driven Macrophage Signature” (LDMS) based on genes downregulated by the p300 inhibitor C646 in PDAC-CM-treated macrophages. In the TCGA-PAAD cohort, Kaplan-Meier analysis demonstrated that patients in the high-LDMS group exhibited shorter overall survival compared to the low-LDMS group (Figure 7A). Furthermore, elevated LDMS scores positively correlated with advanced TNM stages (Figure 7B), linking macrophage lactylation to disease progression.

**Figure 7.**
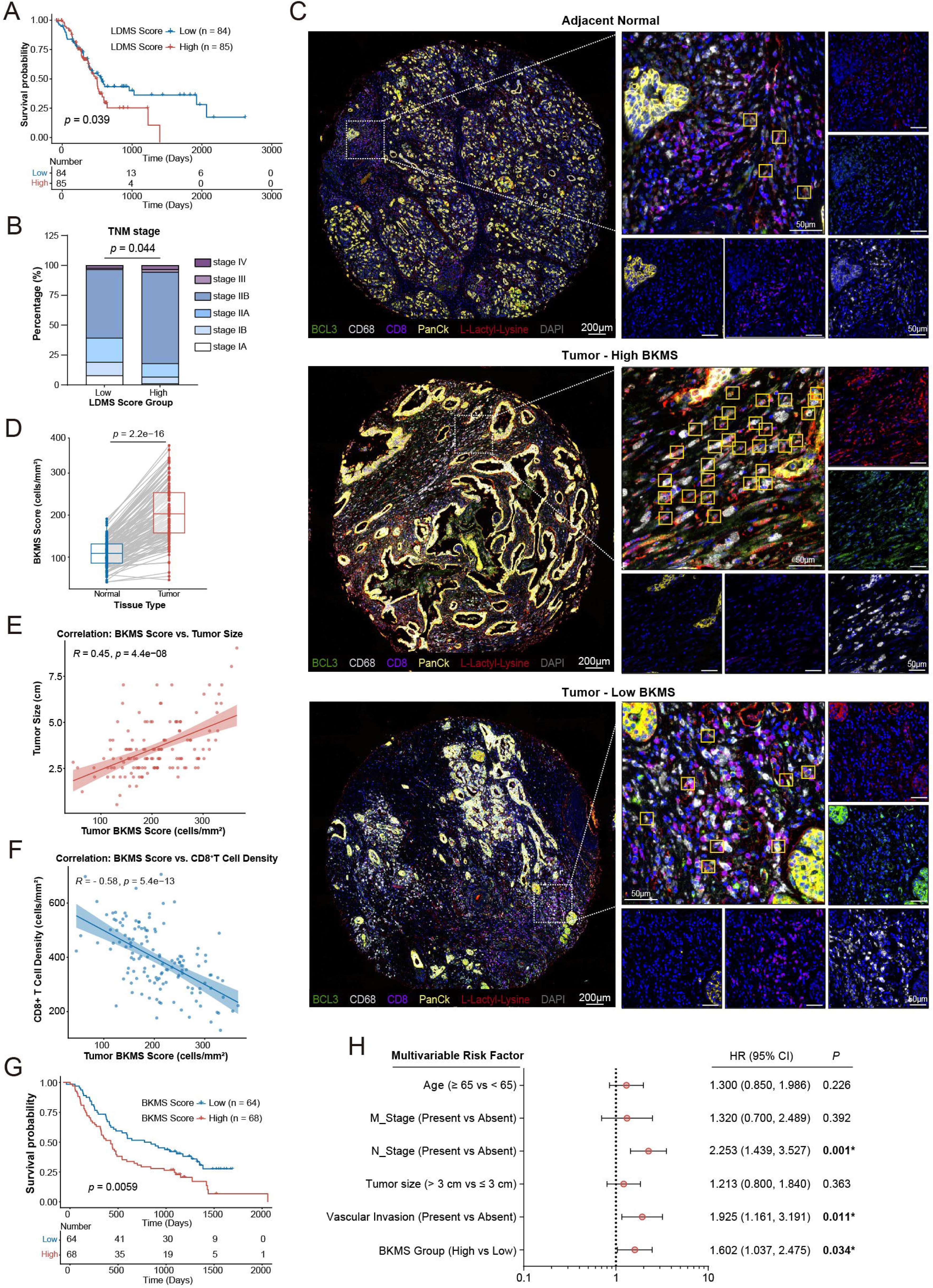
Lactylation-driven macrophage reprogramming correlates with poor prognosis and immune exclusion in PDAC. **(A)** Kaplan-Meier survival analysis of overall survival (OS) in the TCGA-PAAD cohort stratified by the LDMS score (high vs. low; Fleming-Harrington test). **(B)** Distribution of LDMS scores across different TNM stages in the TCGA-PAAD cohort. **(C)** Representative mIF images of human PDAC TMA stained for CD68 (macrophages), BCL3, Pan-Kla, CD8 (cytotoxic T cells), and CK19 (tumor cells). **(D)** Comparison of BKMS between paired tumor and adjacent normal tissues in the TMA cohort. **(E)** Correlation between BKMS and primary tumor size. **(F)** Correlation between BKMS and CD8^+^ T cell density in PDAC tissues. **(G)** Kaplan-Meier curves for OS of patients in the TMA cohort, stratified by BKMS (high vs. low). **(H)** Forest plot displaying hazard ratios (HR) and 95% confidence intervals (CI) from multivariate Cox proportional hazards regression analysis of BKMS and clinicopathological factors.

To confirm the *in situ* involvement of the lactate-macrophage-BCL3 axis, we analyzed a tissue microarray (TMA) of paired PDAC samples using mIF for Pan-Kla, CD68, BCL3, CD8, and CK19 (Figure 7C). Representative images illustrate the spatial distribution of these markers across adjacent normal tissues and PDAC tissues, grouped by low or high BCL3-Kla Macrophage Scores (BKMS) (Figure 7C). Quantitative analysis confirmed that the BKMS was significantly elevated in tumor tissues compared to matched adjacent normal tissues (Figure 7D). To assess the relationship between macrophage lactylation and tumor burden, we analyzed clinical characteristics and observed a positive correlation between BKMS and primary tumor size (Figure 7E). Crucially, we observed an inverse correlation between BKMS and cytotoxic T cell infiltration. Tumor regions rich in BCL3-lactylated macrophages displayed a marked deficiency of CD8^+^ T cells, representing an immune-excluded phenotype (Figure 7F). Clinically, elevated BKMS was associated with significantly poorer overall survival (Figure 7G). Multivariate Cox regression analysis further identified the BKMS as an independent prognostic predictor for PDAC (Figure 7H).

Next, we investigated the immunological consequences of macrophage lactylation by mapping the TME landscape against the LDMS in the TCGA cohort. ESTIMATE analysis indicated that high-LDMS tumors exhibited significantly higher Immune and Stromal Scores (Figure S7A), suggesting a stroma-rich microenvironment. Deconvolution via CIBERSORT revealed that the LDMS score was robustly correlated with the infiltration of M2 macrophages (Figure S7B). Notably, scatter plot analysis highlighted a stronger positive correlation between LDMS and M2 macrophages in the high-score group compared to the low-score group (Figure S7C), suggesting a functional coupling between lactylation intensity and M2 polarization. Further functional characterization using GSVA demonstrated that the LDMS score was highly correlated with signatures of immune checkpoints, Tregs, and T cell co-inhibition, while showing weaker associations with markers of cytolytic activity (Figure S7D). A bidirectional scatter plot mapping T cell co-stimulation versus Checkpoint signatures further confirmed that high-LDMS tumors preferentially clustered in the quadrant defined by high Checkpoint expression (Figure S7E). Collectively, these analyses indicate that the BCL3-lactylation axis is associated with a highly immunosuppressive, M2-dominant microenvironment, which correlates with the spatial exclusion of cytotoxic T cells and poor clinical outcomes.

## Discussion

Metabolic reprogramming fuels malignancy, yet the mechanistic links between aberrant glycolysis and the pro-tumorigenic microenvironment remain incompletely defined.^31^ Our study identifies a critical metabolic-signaling axis linking cancer cell glycolysis to TAMs reprogramming in PDAC. We demonstrate that tumor-derived lactate acts not merely as a metabolic by-product but as a direct signaling molecule driving macrophage reprogramming. Specifically, L-lactylation of the transcriptional cofactor BCL3 at K21 serves as the central trigger for TAMs remodeling. Mechanistically, the PTM stabilizes the interaction between BCL3 and NF-κB p50 homodimers, thereby competitively displacing pro-inflammatory p65 subunits. The resulting transcriptional shift drives macrophages to support tumor cell proliferation via direct signaling, creating a niche that facilitates malignant progression.

Our study broadens the current understanding of the TME crosstalk by defining a functional “lactate shuttle” between cancer cells and macrophages. Unlike prior studies focusing on lactate primarily as a carbon source or acidification agent,^32,33^ we demonstrate an active transport mechanism where lactate is exported and imported via highly specific transporters. The metabolic interplay prompts a re-evaluation of macrophages not merely as immune effectors, but as metabolic gatekeepers.

By scavenging tumor-derived lactate, TAMs not only function as a metabolic buffer that protects cancer cells from local acidotoxicity, but also utilize the imported metabolite to acquire a tumor-promoting phenotype. The metabolic interplay highlights that PDAC progression relies on a co-evolved ecosystem where macrophages couple metabolic clearance with phenotype remodeling. Our data explicitly distinguish the effects of extracellular lactate as a reprogramming substrate from those of intracellular self-generated lactate. The failure of acetate and D-lactate to recapitulate the phenotypic remodeling observed with L-lactate confirms that macrophage polarization relies on an enzymatic, stereoselective sensing mechanism rather than a generalized stress response to acidification.

A critical finding of our study is the multidimensional effect of lactate-reprogrammed TAMs in promoting PDAC progression. Our *in vitro* co-culture assays demonstrated that Lac-PrM significantly accelerated PDAC cell proliferation even in the absence of adaptive immune cells, indicating a direct paracrine contribution to malignant growth. While our functional validation primarily highlights the direct trophic support, the broad upregulation of M2-associated genes like ARG1 and IL10 strongly implicates a potential concurrent role in immune exclusion. As representative examples of the immunosuppressive program, high Arginase-1 (ARG1) expression depletes the microenvironmental arginine pool to metabolically arrest T cell proliferation.^34,35^ Similarly, IL-10 acts as a potent cytokine to dampen cytotoxic responses.^36,37^ Bioinformatics analysis of the TCGA cohort and spatial analysis of TMA further support the immune-dependent contribution by revealing signatures of T cell exhaustion and the spatial exclusion of CD8^+^ T cells in regions enriched with BCL3-lactylated macrophages. The inverse correlation between BCL3-lactylated macrophages and CD8^+^ T cell infiltration highlights the potential of lactylated macrophages to act as a biochemical barrier arresting T cells before reaching the tumor parenchyma. Together, the correlative data suggest that the BCL3-lactylation axis drives malignant progression primarily through paracrine signaling, with a highly probable concurrent contribution from immune evasion.

Furthermore, our findings broaden the functional landscape of the “lactylome” beyond chromatin regulation in PDAC-associated macrophages. Since the initial discovery of histone lactylation (e.g., H3K18la) as an epigenetic regulator,^38,39^ non-histone targets have remained underexplored.^40^ The evidence presented here indicates that lactylation extends beyond chromatin regulation to directly modulate signal transduction. Distinct from histone modifications that shape the stable epigenetic landscape, BCL3 lactylation likely induces rapid conformational changes that alter protein-protein interaction interfaces, thereby allowing macrophages to respond dynamically to metabolic fluctuations within the TME.

At the molecular level, our data clarify the specific role of BCL3 in NF-κB signaling within PDAC-associated macrophages. While BCL3 is classically described as either a co-activator or co-repressor depending on its binding partners,^17^ our results demonstrate that K21 lactylation dictates its repressor function by stabilizing the BCL3-p50 homodimer complex. Structural modeling and functional assays indicate that L-lactylation at K21 induces conformational remodeling of the N-terminus to expose the NLS, driving nuclear entry and subsequent ARD-mediated binding. By sequestering p50, lactylated BCL3 blocks p65 binding and attenuates the canonical NF-κB pathway.^41^ The resulting transcriptional landscape explains why TAMs remain functionally quiescent regarding anti-tumor immunity (loss of p65 activity) while actively supporting tumor growth (gain of p50/BCL3-driven M2 functions), effectively creating a permissive niche that fosters immune tolerance.^42-44^

Several study limitations warrant consideration. First, while pharmacologic inhibition and site-specific mutagenesis validated BCL3 lactylation function, specific “writer” (lactate transferase) and “eraser” (delactylase) enzymes regulating K21 remain to be fully characterized. However, identification of the specific modification site (K21) and its structural impact on BCL3-p50 interaction provides a sufficient mechanistic basis for current findings, laying the groundwork for future enzymatic screens. Second, although C646 inhibits both acetylation and lactylation, consistent phenotypic mimicry between C646 treatment and genetic K21R mutants strongly supports lactylation as the dominant driver in this context, though future development of specific chemical probes remains necessary. Finally, while robust in murine models and patient tissues, temporal dynamics of lactylation during early PDAC stages require longitudinal evaluation. Nevertheless, the strong correlation established here highlights the potential of BCL3-K21la as a stratification biomarker for metabolic-immune targeted therapies.

In conclusion, our findings unveil a metabolic-signaling axis wherein PDAC-derived lactate promotes a pro-tumorigenic macrophage phenotype via BCL3-K21 lactylation. Such mechanism reveals how the hyper-glycolytic PDAC feature is mechanistically coupled to macrophage reprogramming. By exploiting the BCL3-p50 axis via site-specific lactylation, PDAC cells effectively re-educate macrophages to serve as obligate partners in malignant progression. Targeting the BCL3 L-lactylation axis offers a promising avenue to disrupt the metabolic dependency, restoring the anti-tumor potential of the innate immune system while depriving the tumor of essential macrophage-mediated support.

## Contributions

Conceptualization: Y.-W.S., D.-J.L., S.-H.J., Y.-J.H., and P.-X.J.; Methodology: P.-X.J., J.-H.Z., X.-S.M., F.Y., and Y.-W.L.; Software and Formal Analysis: X.-S.M., F.Y., and J.- Y.Y.; Investigation: P.-X.J., J.-H.Z., X.-S.M., F.Y., Y.-W.L., Q.W.,and K.X.; Resources: W.L., X.-L.F., Y.-M.H., W.-Y.G., and Y.-J.H.; Data Curation: P.-X.J., J.-H.Z., and X.-S.M.; Writing-Original Draft: P.-X.J., J.-H.Z., and D.-J.L.; Writing-Review & Editing: Y.-W.S., D.-J.L., S.-H.J., Y.-J.H., P.-X.J., and J.-H.Z.; Visualization: P.-X.J., X.-S.M., and F.Y.; Supervision: Y.-W.S., D.-J.L., S.-H.J., and Y.-J.H.; Project Administration: Y.-W.S. and D.-J.L.; Funding Acquisition: Y.-W.S., J.-Y.Y.,W.L., M.-W.Y. and Y.-M.H

## Financial support statement and Acknowledgement

This study was supported by the National Natural Science Foundation of China (82472917 to Y.-W.S., 82473029 to J.-Y.Y., 82573865 to W.L.), the Bethune Charitable Foundation (YXZL-2023-B-002 of Y.-M.H) and the Natural Science Foundation of Shanghai (24ZR1445700 of M.-W.Y.).

