## Supplementary Material for "L-Lactate reprograms tumor-associated macrophages to drive pancreatic cancer progression via BCL3 lactylation"

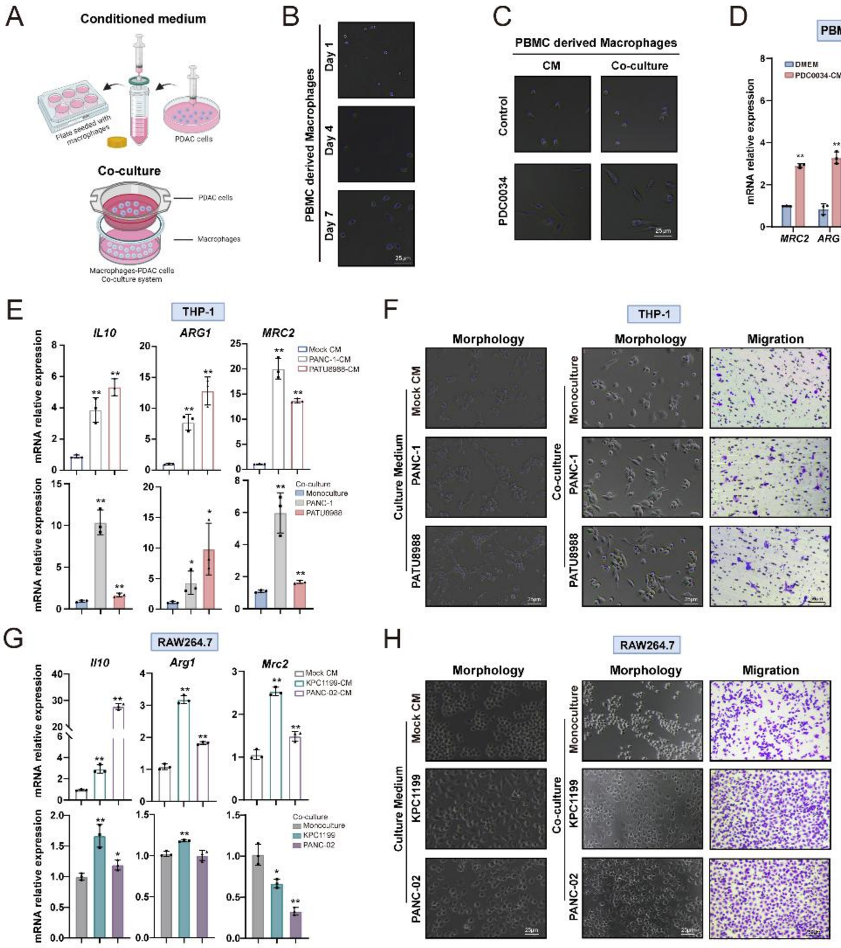


**Figure S1. PDAC-derived signals induce morphological and phenotypic transition in macrophages *in vitro*. (A)** Schematic illustration of the macrophage polarization assays, involving either direct co-culture with PDAC cells or stimulation with PDAC-CM for 24 h. **(B-C)** Representative bright-field microscopy images displaying the differentiation of PBMC-derived monocytes into macrophages **(B)** and their morphological transition following co-culture with primary PDAC cells **(C). (D)** Relative mRNA levels of M2-related genes in human PBMC-derived macrophages following co-culture with primary PDAC cells (PDC0034) or exposure to PDAC-CM (mean ± SEM, unpaired two-tailed *t* test, n = 3 per group). **(E-H)** Relative mRNA levels of M2-related genes, morphology and migration ability in human THP-1 **(E-F)** and murine RAW264.7 macrophages **(G-H)** following co-culture with the indicated PDAC cell lines or exposure to their respective CM (mean ± SEM, unpaired two-tailed *t* test, n = 3 per group). *$P < 0.05$; **$P < 0.01$.

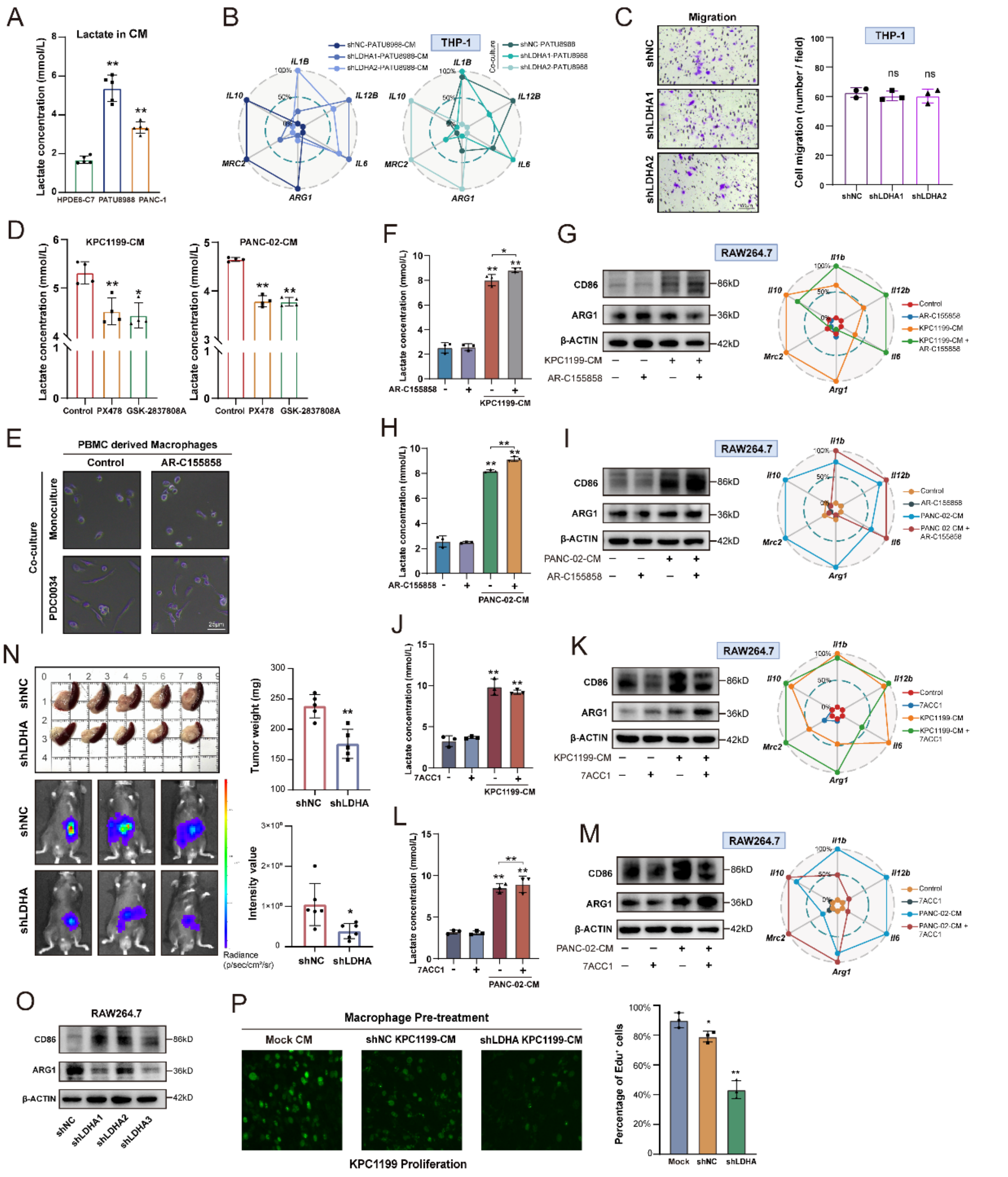


**Figure S2. Targeting lactate transport dynamics impairs M2-like macrophage polarization. (A)** Quantification of lactate concentrations in the culture supernatants of various PDAC cell lines (mean ± SEM, unpaired two-tailed *t* test, n = 5 per group). **(B)** qPCR analysis of polarization markers in THP-1 macrophages co-cultured with control or *LDHA*-deficient tumor cells. **(C)** Transwell migration assays performed on THP-1 cells stimulated with CM derived from control or *LDHA*-KD PATU8988 cells (mean ± SEM, unpaired two-tailed *t* test, n = 3 per group). **(D)** Assessment of supernatant lactate levels in KPC1199 and PANC-02 cells treated with LDH inhibitors (PX-478 or GSK-2837808A)

(mean ± SEM, unpaired two-tailed *t* test, n = 4 per group). **(E)** Representative bright-field microscopy images displaying morphological alterations of PBMC-derived macrophages co-cultured with PDC0034 cells ± AR-C155858. **(F-I)** Evaluation of macrophage function upon lactate influx inhibition. RAW264.7 cells were stimulated with CM from KPC1199 **(F, G)** or PANC-02 **(H, I)** ± AR-C155858. Assays included quantification of intracellular lactate, immunoblot analysis of ARG1/CD86, and qPCR profiling of immune-related genes. **(J-M)** Assessment of macrophage function upon lactate efflux inhibition. RAW264.7 cells were stimulated with CM from KPC1199 **(J, K)** or PANC-02 **(L, M)** in the presence of 7ACC1, followed by measurement of intracellular lactate accumulation and expression of polarization markers. **(N)** Assessment of tumor burden via bioluminescence imaging and terminal tumor weight (mean ± SEM, unpaired two-tailed *t* test, n = 5 per group). **(O)** Immunoblot analysis of polarization markers in RAW264.7 cells cultured with supernatants collected from *Ldha*-KD KPC1199 cells. **(P)** Representative EdU images and quantitative proliferation assays of KPC1199 cells cultured with secondary CM derived from the pre-treated RAW264.7 macrophages (mean ± SEM, unpaired two-tailed *t* test, n = 3 per group). *$P$ < 0.05; **$P$ < 0.01.

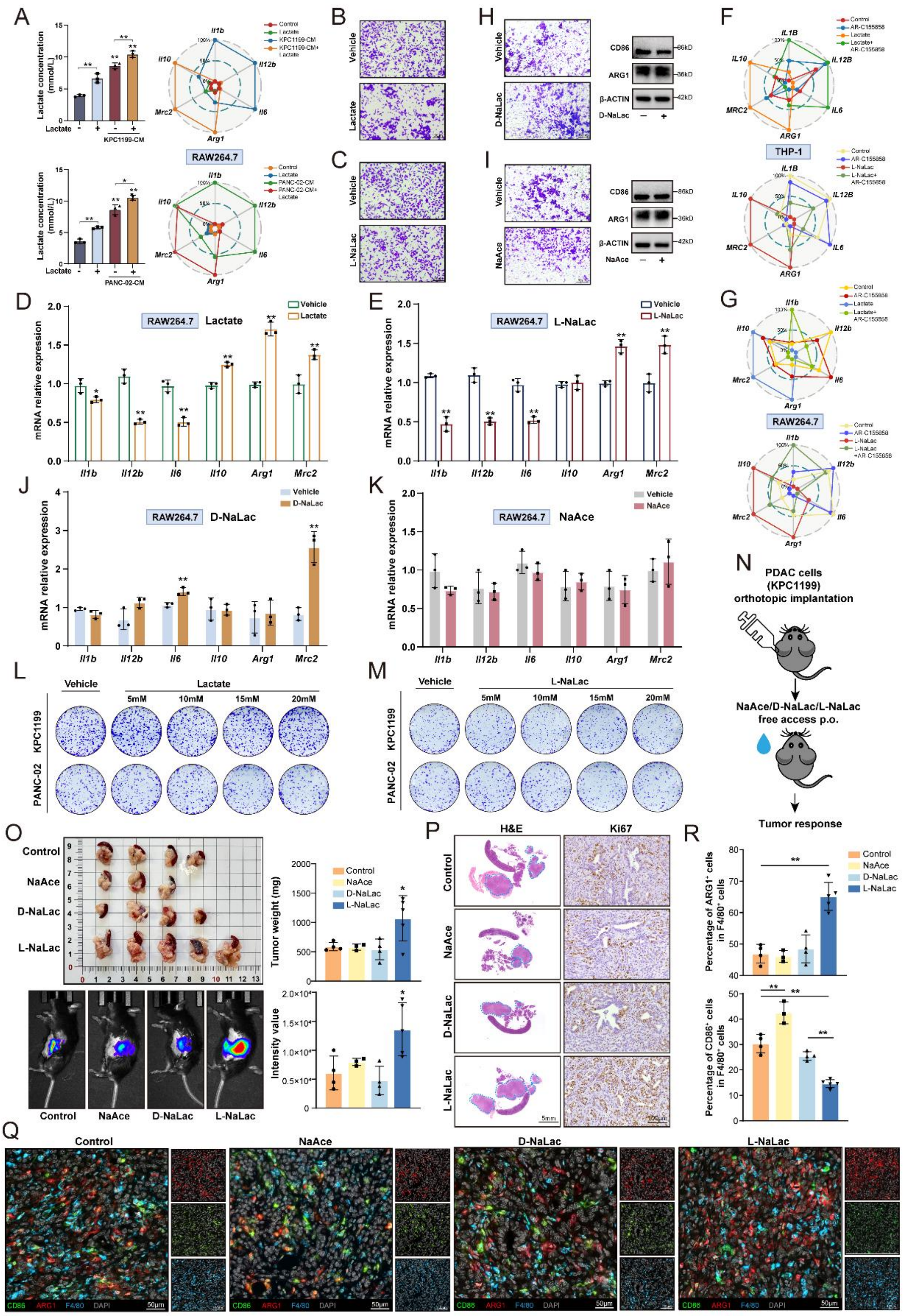


**Figure S3. Specificity of L-lactate in driving tumor-promoting functions of macrophages. (A)** Quantification of intracellular lactate accumulation and gene expression profiles in RAW264.7 cells stimulated with KPC1199 or PANC-02 CM in the presence or absence of additional lactate supplementation (mean ± SEM, one-way ANOVA with Tukey's test, n = 3 per group). **(B-C)** Assessment of migration in RAW264.7 macrophages exposed to different concentrations of lactate **(B)** or L-NaLac **(C)**. **(D-E)** Relative mRNA expression of polarization markers in RAW264.7 cells stimulated with lactate **(D)** or L-NaLac **(E)** (mean ± SEM, unpaired two-tailed *t* test, n = 3 per group). **(F-G)** Radar plots summarizing the relative mRNA expression of immuno-associated genes in THP-1 **(F)** and RAW264.7 **(G)** cells treated with lactate or L-sodium lactate in the presence or absence of AR-C155858. **(H-K)** Evaluation of stereoisomer and metabolite specificity *in vitro*. RAW264.7 cells were treated with D-NaLac **(H, J)** or NaAce **(I, K)**. Panels display migration and immunoblotting of ARG1/CD86 **(H, I)**, alongside immune gene profiling **(J, K)** (mean ± SEM, unpaired two-tailed *t* test, n = 3 per group). **(L-M)** Colony formation assays assessing the direct effect of lactate **(L)** or L-NaLac **(M)** on the proliferation of KPC1199 and PANC-02 cells. **(N)** Schematic illustration of the orthotopic PDAC mouse model treated with vehicle, NaAce, D-NaLac, or L-NaLac via drinking water. **(O)** Tumor burden assessment via bioluminescence imaging and terminal tumor weights (mean ± SEM, one-way ANOVA with Tukey's test, n = 4 for Control, n = 3 for NaAce, n = 4 for D-NaLac, n = 5 for L-NaLac). **(P)** Representative H&E staining, and IHC for Ki67 in PDAC tissue sections. (**Q-R**) Representative mIF staining **(Q)** and quantification **(R)** comparing the $ARG1^{+}$ / $CD86^{+}$ ratio (mean ± SEM, one-way ANOVA with Tukey's test). *$P < 0.05$; **$P < 0.01$.

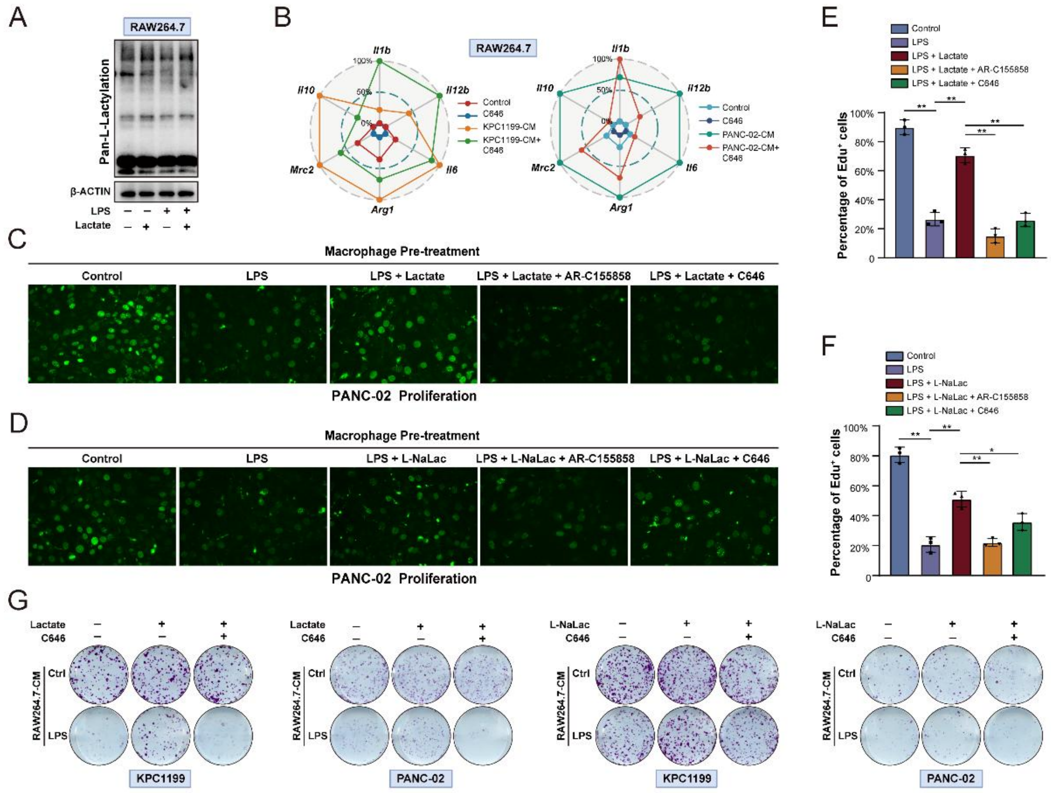


**Figure S4. L-Lactylation is essential for macrophage-supported tumor proliferation.** **(A)** Immunoblot analysis of global L-lactylation levels in RAW264.7 cells stimulated with LPS (inducing endogenous glycolysis) or exogenous lactate. **(B)** Radar plots profiling immune-related gene expression in RAW264.7 cells stimulated with PDAC-CM in the presence or absence of C646. **(C-F)** Representative EdU staining **(C, D)** and quantification **(E, F)** of PANC-02 proliferation induced by CM from macrophages stimulated with LPS, lactate or L-sodium lactate, in the presence or absence of AR-C155858 or C646 (mean ± SEM, one-way ANOVA with Tukey's test, n = 3 per group). **(G)** Colony formation assays assessing the proliferation of PDAC cells cultured with CM from macrophages treated with lactate or L-sodium lactate, with or without C646 co-treatment. *$P < 0.05$; **$P < 0.01$.

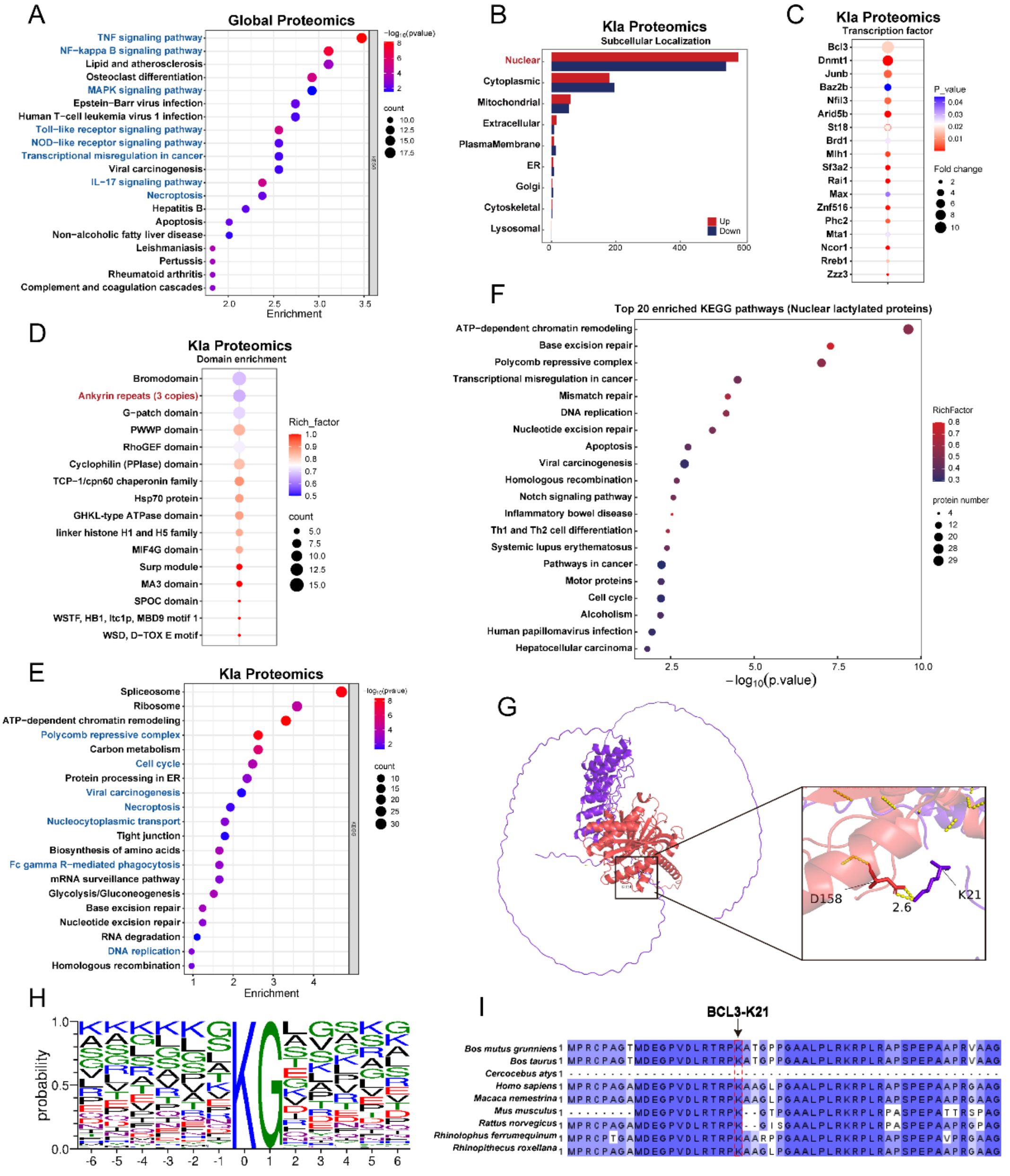


**Figure S5. Bioinformatic characterization of the macrophage L-lactylome. (A)** KEGG pathway enrichment analysis of differentially expressed proteins in the global proteome (CM-treated vs. Control). **(B)** Subcellular localization analysis of differentially L-lactylated proteins. **(C)** Heatmap visualizing the enrichment of differentially lactylated transcription factors. **(D)** Protein domain enrichment analysis of the differentially lactylated proteome. **(E)** KEGG pathway enrichment analysis of differentially lactylated proteins. **(F)** KEGG pathway enrichment analysis specifically performed on differentially lactylated nuclear proteins. **(G)** Structural modeling of the P300 HAT domain (Red) and BCL3 (Purple) complex. Left: Global interaction view. Right: Zoom-in view showing the predicted hydrogen bond (2.6 Å) between BCL3-K21 and P300-D158. **(H)** Sequence motif analysis centered on the K21 residue. **(I)** Evolutionary conservation analysis of the BCL3 K21 residue across diverse species.

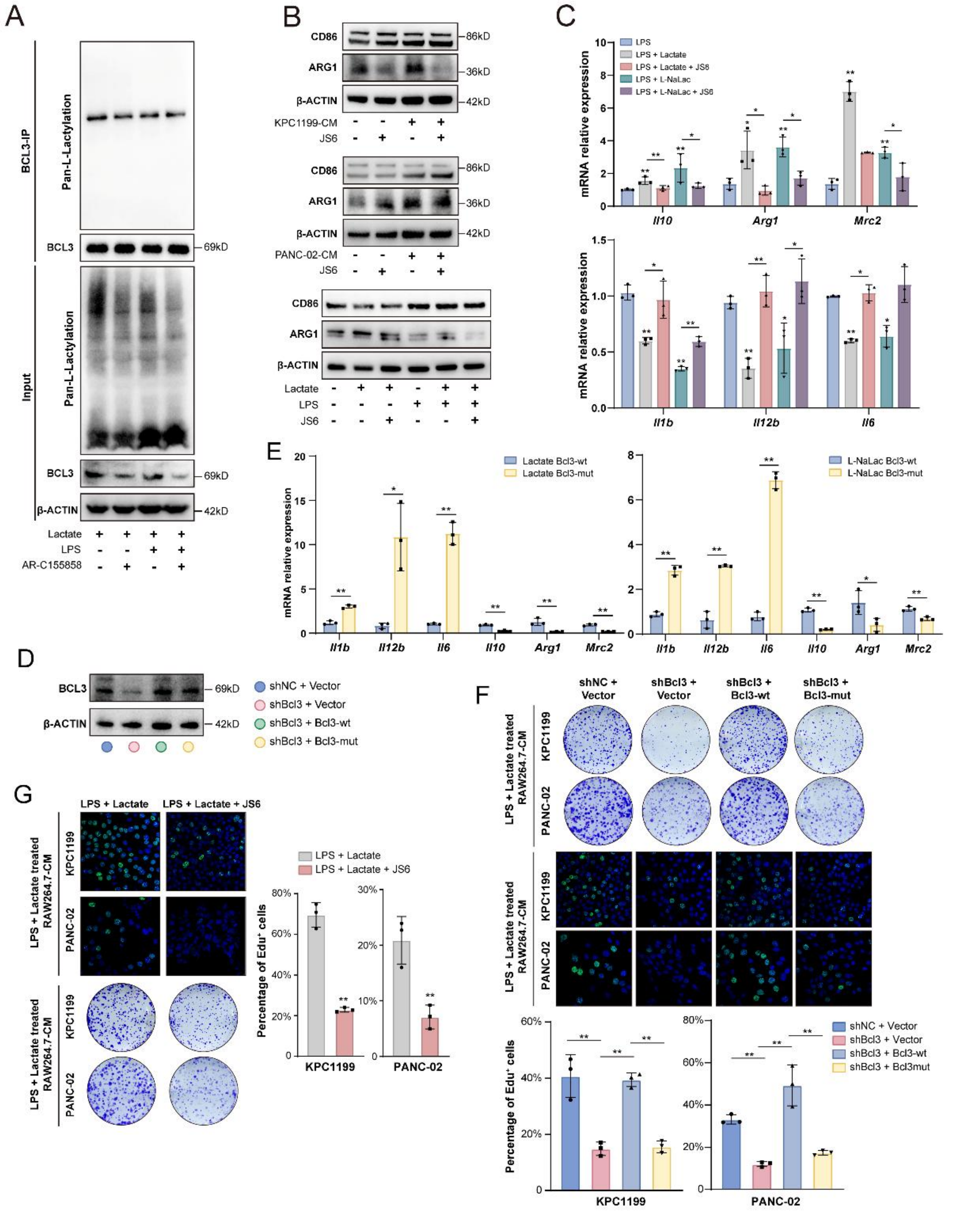


**Figure S6. Validation of the Lactate-BCL3-NF-κB axis and K21R functional defects.** **(A)** Co-IP assays assessing BCL3 L-lactylation levels (Pan-Kla) in RAW264.7 cells stimulated with lactate ± the MCT1/2 inhibitor AR-C155858**.** **(B-C)** Functional validation of the BCL3-NF-κB axis. **(B)** Immunoblotting of ARG1 and CD86 and **(C)** qPCR analysis of immune-associated genes in RAW264.7 cells stimulated with PDAC-CM or LPS/Lactate ± JS6 (mean ± SEM, one-way ANOVA with Tukey's test, n = 3). **(D)** Immunoblot confirmation of *Bcl3* KD and rescue efficiency in RAW264.7 cells reconstituted with WT BCL3 or the K21R mutant. **(E)** qPCR analysis of immune-associated genes in RAW264.7 cells expressing WT or K21R mutant BCL3, treated with lactate or L-NaLac (mean ± SEM, unpaired two-tailed *t* test, n = 3 per group). **(F)** Evaluation of PDAC cell proliferation via colony formation assays and EdU staining following culture with CM from lactate-stimulated WT or K21R RAW264.7 cells (mean ± SEM, one-way ANOVA with Tukey's test, n = 3). **(G)** Representative EdU staining, quantification, and colony formation images of KPC1199 and PANC-02 cells cultured with CM from RAW264.7 cells ± JS6 (mean ± SEM, unpaired two-tailed *t* test, n = 3). *$P < 0.05$; **$P < 0.01$.

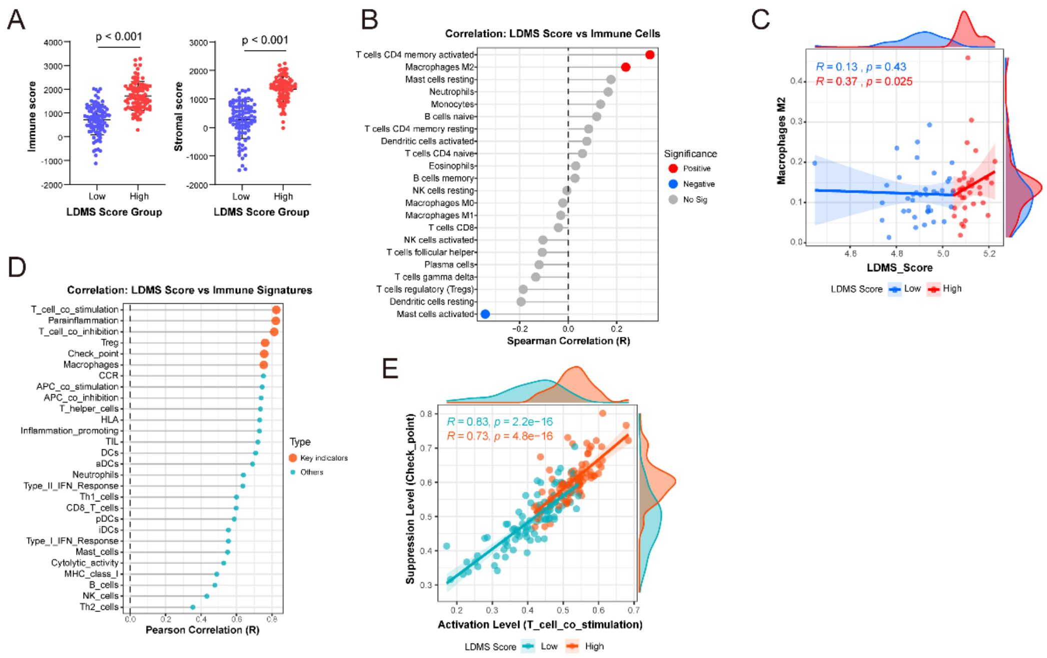


**Figure S7. The LDMS defines an immunosuppressive and exhausted TME landscape.** **(A)** Comparison of ESTIMATE Immune and Stromal Scores between high- and low-LDMS groups in the TCGA-PAAD cohort. **(B)** Lollipop chart showing the correlation between LDMS scores and the abundance of 22 immune cell types (CIBERSORT). Note the specific positive correlation with M2 macrophages. **(C)** Scatter plots comparing the correlation strength between LDMS scores and M2 macrophage abundance in high- versus low-LDMS groups. **(D)** Lollipop chart displaying the correlation between LDMS scores and specific immune functional signatures (GSVA), highlighting the enrichment of inhibitory and exhaustion-related pathways. **(E)** Bidirectional scatter plot visualizing the clustering of high-LDMS tumors in the quadrant defined by high Checkpoint and high T cell co-stimulation signatures.

**Supplementary Table S1**

| Antibodies | | | |
|---|---|---|---|
| **Name** | **Company** | **Catalogue** | **RRID** |
| Pan-Keratin Rabbit Monoclonal Antibody | Cell Signaling Technology (Danvers, MA, USA) | 83957 | / |
| Cytokeratin 19 Polyclonal antibody | Proteintech Group (Wuhan, China) | 10712-1-AP | AB_2133325 |
| CD8a Monoclonal antibody | Proteintech Group (Wuhan, China) | 66868-1-Ig | AB_2882205 |
| Anti-CD68 antibody | Abcam (Cambridge, UK) | ab213363 | AB_2801637 |
| CD68 Rabbit Monoclonal Antibody | Cell Signaling Technology (Danvers, MA, USA) | 97778 | / |
| CD16 Polyclonal antibody | Proteintech Group (Wuhan, China) | 16559-1-AP | AB_2878279 |
| Arginase-1 Polyclonal antibody | Proteintech Group (Wuhan, China) | 16001-1-AP | AB_2289842 |
| CD86 Rabbit Monoclonal Antibody | Cell Signaling Technology (Danvers, MA, USA) | 91882 | AB_2797422 |
| beta-actin antibody | Share-bio (Shanghai, China) | SB-AB0035 | AB_3674188 |
| LDHA-Specific Polyclonal antibody | Proteintech Group (Wuhan, China) | 19987-1-AP | AB_10646429 |
| F4/80 Polyclonal antibody | Proteintech Group (Wuhan, China) | 28463-1-AP | AB_2881149 |
| Ki-67 Polyclonal antibody | Proteintech Group (Wuhan, China) | 27309-1-AP | AB_2756525 |
| Anti-L-Lactyl Lysine Rabbit mAb | PTM Biolabs (Chicago, IL, USA) | PTM-1401RM | AB_2942013 |
| Anti-Acetyllysine Mouse mAb | PTM Biolabs (Chicago, IL, USA) | PTM-101 | AB_2940830 |

| | | | |
|---|---|---|---|
| BCL3 Rabbit pAb | Zen Bioscience (Chengdu, China) | 860954 | / |
| Anti-NF-kappaB p105/p50 Rabbit Polyclonal Antibody | Genuin Biotech (Hangzhou, China) | 61524 | / |
| NF-κB p65 Recombinant antibody | Proteintech Group (Wuhan, China) | 80979-1-RR | AB_2918923 |
| KD-Validated Mannose Receptor (CD206) Recombinant Rabbit mAb | Genuin Biotech (Hangzhou, China) | 61279 | / |
| Histone H3 Polyclonal antibody | Proteintech Group (Wuhan, China) | 17168-1-AP | AB_2716755 |
| Goat Anti-Rabbit IgG (H+L) HRP | Share-bio (Shanghai, China) | SB-AB0101 | / |
| Goat Anti-Mouse IgG (H+L) HRP | Share-bio (Shanghai, China) | SB-AB0102 | / |
| Cy3-Conjugated Goat Anti-Rabbit IgG (H&L) Secondary Antibody | Genuin Biotech (Hangzhou, China) | 324 | / |

### Supplementary Table S2

| Target Sequence | | |
|---|---|---|
| **Name** | **Species** | **Catalogue** |
| *IL6* | Human | Forward: ACTCACCTCTTCAGAACGAATTG<br>Reverse: CCATCTTTGGAAGGTTCAGGTTG |
| *IL12B* | Human | Forward: ACCCTGACCATCCAAGTCAAA<br>Reverse: TTGGCCTCGCATCTTAGAAAG |
| *IL1B* | Human | Forward: ATGATGGCTTATTACAGTGGCAA<br>Reverse: GTCGGAGATTCGTAGCTGGA |
| *MRC2* | Human | Forward: CCGAAACCGGCTATTCAACCT<br>Reverse: CGGTCACACTCATACATGCCC |
| *ARG1* | Human | Forward: GTGGAAACTTGCATGGACAAC<br>Reverse: AATCCTGGCACATCGGGAATC |
| *IL10* | Human | Forward: GACTTTAAGGGTTACCTGGGTTG<br>Reverse: TCACATGCGCCTTGATGTCTG |
| *SLC16A1* | Human | Forward: AGGTCCAGTTGGATACACCCC<br>Reverse: GCATAAGAGAAGCCGATGGAAAT |
| *SLC16A7* | Human | Forward: GGGTTGGATTGTGGTTGGAG<br>Reverse: TCCTGCGTACATAACAGCCAG |
| *SLC16A3* | Human | Forward: CCATGCTCTACGGGACAGG<br>Reverse: GCTTGCTGAAGTAGCGGTT |
| *Il6* | Murine | Forward: TAGTCCTTCCTACCCCAATTTCC<br>Reverse: TTGGTCCTTAGCCACTCCTTC |
| *Il12b* | Murine | Forward: TGGTTTGCCATCGTTTTGCTG<br>Reverse: ACAGGTGAGGTTCACTGTTTCT |

| | | |
|---|---|---|
| *Il1b* | Murine | Forward: GCAACTGTTCCTGAACTCAACT<br>Reverse: ATCTTTTGGGGTCCGTCAACT |
| *Mrc2* | Murine | Forward: TCTCCCGGAACCGACTCTTC<br>Reverse: GGTCGAGCACATAGGTCTTCT |
| *Arg1* | Murine | Forward: CTCCAAGCCAAAGTCCTTAGAG<br>Reverse: AGGAGCTGTCATTAGGGACATC |
| *Il10* | Murine | Forward: GCTCTTACTGACTGGCATGAG<br>Reverse: CGCAGCTCTAGGAGCATGTG |
| sh-*LDHA* | Human | #1: CCACCATGATTAAGGGTCTTT<br>#2: CCAAAGATTGTCTCTGGCAAA |
| sh-*Ldha* | Murine | #1: CCTTCCTGTGGTGTGCTTCAT<br>#2: GCCGAGAGCATAATGAAGAAT |
| sh-*Bcl3* | Murine | OmicsLink™ shRNA Expression Clone (Cat# MSH036845-LVRU6GP-a; GeneCopoeia, Inc.) |
| OE-*Bcl3*(WT) | Murine | OmicsLink™ Expression Clone encoding wild-type murine *Bcl3* (Cat# CS-Mm24400-Lv158-01; GeneCopoeia, Inc., Rockville, MD). |
| OE-*Bcl3*(K21R) | Murine | OmicsLink™ Expression Clone encoding lactylation-deficient K21R mutant murine *Bcl3* (Cat# CS-Mm24400-Lv158-02; GeneCopoeia, Inc., Rockville, MD). |
